# Epigenetic Memory *via* Concordant DNA Methylation is Inversely Correlated to Developmental Potential of Mammalian Cells

**DOI:** 10.1101/072488

**Authors:** Minseung Choi, Diane P. Genereux, Jamie Goodson, Haneen Al-Azzawi, Shannon Q. Allain, Noah Simon, Stan Palasek, Carol B. Ware, Chris Cavanaugh, Daniel G. Miller, Winslow C. Johnson, Kevin D. Sinclair, Reinhard Stöger, Charles D. Laird

## Abstract

In storing and transmitting epigenetic information, organisms must balance the need to maintain information about past conditions with the capacity to respond to information in their current and future environments. Some of this information is encoded by DNA methylation, which can be transmitted with variable fidelity from parent to daughter strand. High fidelity confers strong pattern matching between the strands of individual DNA molecules and thus pattern stability over rounds of DNA replication; lower fidelity confers reduced pattern matching, and thus greater flexibility. • Here, we present a new conceptual framework, Ratio of Concordance Preference (RCP), that uses double-stranded methylation data to quantify the flexibility and stability of the system that gave rise to a given set of patterns. • We find that differentiated mammalian cells operate with high DNA methylation stability, consistent with earlier reports. Stem cells in culture and in embryos, in contrast, operate with reduced, albeit significant, methylation stability. We conclude that preference for concordant DNA methylation is a consistent mode of information transfer, and thus provides epigenetic stability across cell divisions, even in stem cells and those undergoing developmental transitions. Broader application of our RCP framework will permit comparison of epigenetic-information systems across cells, developmental stages, and organisms whose methylation machineries differ substantially or are not yet well understood.

## Introduction

Organismal development is characterized by a shift from the phenotypic flexibility of embryonic cells to the canalized identities of differentiated cells. To achieve stable gene-regulatory states in terminally differentiated cells, organisms ranging from Archaea to humans use a variety of epigenetic mechanisms, including DNA methylation. Perturbation of the state of DNA methylation at various loci in differentiated cells is associated with several human cancers [1–3]. In turn, restoring epigenetic flexibility of some loci has proven challenging in efforts to create induced pluripotent stem (iPS) cells [4]. Together, these findings highlight the importance of shifting ratios of epigenetic flexibility and stability in establishing cellular identity.

There exists an extensive literature documenting changes in single-locus and genome-wide methylation frequencies at various stages of development [5, 6]. Most genomic regions in primordial germ cells (PGCs), for example, are known to undergo dramatic and rapid shifts in DNA methylation frequency [7]. It is now clear that mammalian stem cells can utilize active demethylation [8], highlighting the potential for both gain and loss of cytosine methylation to impact the overall methylation frequency and, perhaps, stability of a given genomic region during development.

High concordance of methylation in differentiated cells, with matching states for parent and daughter DNA strands at individual CpG/CpG dyads, is considered to be a hallmark of conservative epigenetic processes [9–13]. For earlier stages of development, the extent of concordance is far less clear. For example, do methylation patterns in dividing embryonic stem cells arise entirely by random placement of methyl groups, or is concordance favored to some degree?

Recent work has begun to address these issues [7, 14–18]. Shipony *et al.* [16] analyzed single-stranded DNA methylation patterns in populations of cultured cells established from single founder cells. Under this approach, the degree of stability was inferred from the extent of congruence among patterns collected from cultured descendant cells. The observation of substantial pattern diversity among cells separated by many rounds of division led Shipony *et al.* [16] to conclude that the bulk of methylation in human embryonic stem (ES) and induced pluripotent stem (iPS) cells arises through “dynamic” – that is, non-conservative – DNA methylation processes rather than through the “static” – that is, conservative – processes that were emphasized in earlier studies [10, 11, 19]. Using data collected using hairpin-bisulfite PCR [13], which yields double-stranded DNA methylation patterns, other studies suggested that dynamic processes contribute substantially to DNA methylation in cultured mouse ES cells, but perhaps not to the exclusion of the conservative processes that dominate at many loci in adult differentiated cells [7, 14, 15, 17, 18].

To fully characterize the balance between conservative and non-conservative methylation processes, it is necessary to quantify the extent to which the arrangement of methylation in a given set of patterns deviates from the null assumption of random placement. To assess and visualize such deviations, we here introduce a new metric, Ratio of Concordance Preference (RCP), which utilizes double-stranded methylation data. Here, as previously, we use the term *double-stranded DNA methylation pattern* to refer to the overall pattern of methylation on both top and bottom strands of an individual double-stranded DNA molecule. Double-stranded patterns provide information on the extent of matching between methylation states on parent and daughter strands, which are separated by exactly one round of DNA replication. RCP requires no assumptions about the enzymatic mechanisms of methylation and demethylation, and so enables comparison across diverse species and developmental stages.

Jeltsch and Jurkowska [20] have emphasized the balance of methylating and demethylating processes — rather than the propagation of specific methylation patterns — as the primary determinant of the overall set of patterns present in a given cellular population at a given time. In this framework, RCP can be thought of as a metric for quantifying the extent to which the set of patterns produced by a given system of methylating and demethylating processes deviates from the set of patterns expected if methyl groups are placed entirely at random.

In parameterizing RCP, we use the term “conservative”, in lieu of “static” as used previously [16], to describe processes that preferentially establish concordant as opposed to discordant methylation states. We consider non-conservative processes, described previously as “dynamic” [16], as having one of two forms: “random” processes, which add or remove methyl groups with equal preference for concordance and for discordance, and “dispersive” processes, which preferentially establish discordant methylation states.

We validate our RCP framework by confirming its ability to identify systems in which contributions from conservative processes are nearly complete or nearly absent, as well as systems on the continuum between these extremes. We apply this new framework to our authenticated, double-stranded DNA methylation patterns, both published and previously unpublished, collected by dideoxy sequencing from DNA of human and murine cells. To expand the data available for this initial RCP analysis, we also examine double-stranded methylation patterns from three recent publications that used pyrosequencing. [14, 15, 17] Compared to dideoxy sequencing, pyrosequencing can provide greater sequencing depth, but yields considerably shorter reads. To improve our understanding of transitions between stem and differentiated cells, we ask: (*i*) how strong are preferences for concordant DNA methylation states in cultured stem cells?; (*ii*) do concordance preferences change as cultured cells shift between stem and differentiated states?; and (*iii*) in the developing embryo, do stem cells of various potencies have preferences that mirror those in cultured stem cells?

## Results and Discussion

### Ratio of Concordance Preference is Defined for All Possible Configurations of Methylation at Symmetric Nucleotide Motifs

We have developed Ratio of Concordance Preference (RCP) to assess the strategy of binary information transfer, with focus on the degree to which exact information is conserved. We apply our RCP framework to DNA methylation in mammalian cells. Our goal is to infer whether and how much the system of processes that established a given set of methylation patterns prefers concordant to discordant methylation states. This general formulation is free of assumptions about the molecular mechanisms whereby methylation is added to and removed from DNA.

In our data from double-stranded DNA molecules from human and mouse, methylation occurs principally at the CpG motif. This symmetric motif may be written as CpG/CpG, here termed “CpG dyad”. CpG dyads have opportunities for methylation on both strands. The methylation state of a dyad thus takes one of three forms: fully methylated, at frequency *M*, with methylated cytosines on both strands; hemimethylated, at frequency *H*, with a methylated cytosine on only one strand; and unmethylated, at frequency *U*, with neither cytosine methylated. The RCP framework can also be extended to non-CpG methylation at symmetric nucleotide motifs.

To infer concordance preference for sets of double-stranded methylation patterns, we use the overall frequency of methylation, *m*, and the frequency of unmethylated dyads, *U*, of each data set. Because *m* is derived from the three dyad frequencies, the pair (*m*, *U*) encompasses the full information available from the implicit dyad frequencies, *M* and *H*. We evaluate the extent of deviation from expectations under a random model in which the system has no preference for either concordant or discordant placement of methyl groups, using RCP, defined as:

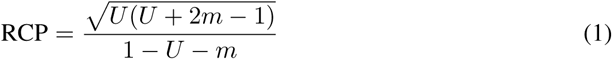

RCP can also be expressed in a form more familiar in biology if dyad frequencies are considered as genotype frequencies for a gene with two alleles. RCP^2^ is 4*M U/H*^2^, which is expected to equal 1 under the Hardy-Weinberg equilibrium [21, 22]. Thus, RCP can be considered as a measure of deviation from random expectations.

The random expectations, for which RCP = 1, are met both with truly random placement of methyl groups, and with equal contributions from processes operating with strong preference for concordance and processes operating with strong preference for discordance. Under this random model, the frequency of unmethylated dyads is given by *U* = (1*-m*)^2^, leading to dyad frequencies as expected under the binomial distribution (Fig 1a,b dashed curve; Fig 1c). A system in which methyl groups are added *de novo* without regard to the methylation state of the other strand [23], such as one dominated by the activity of mammalian Dnmt3s, is expected to behave largely in accordance with random expectations.

**Figure 1.**
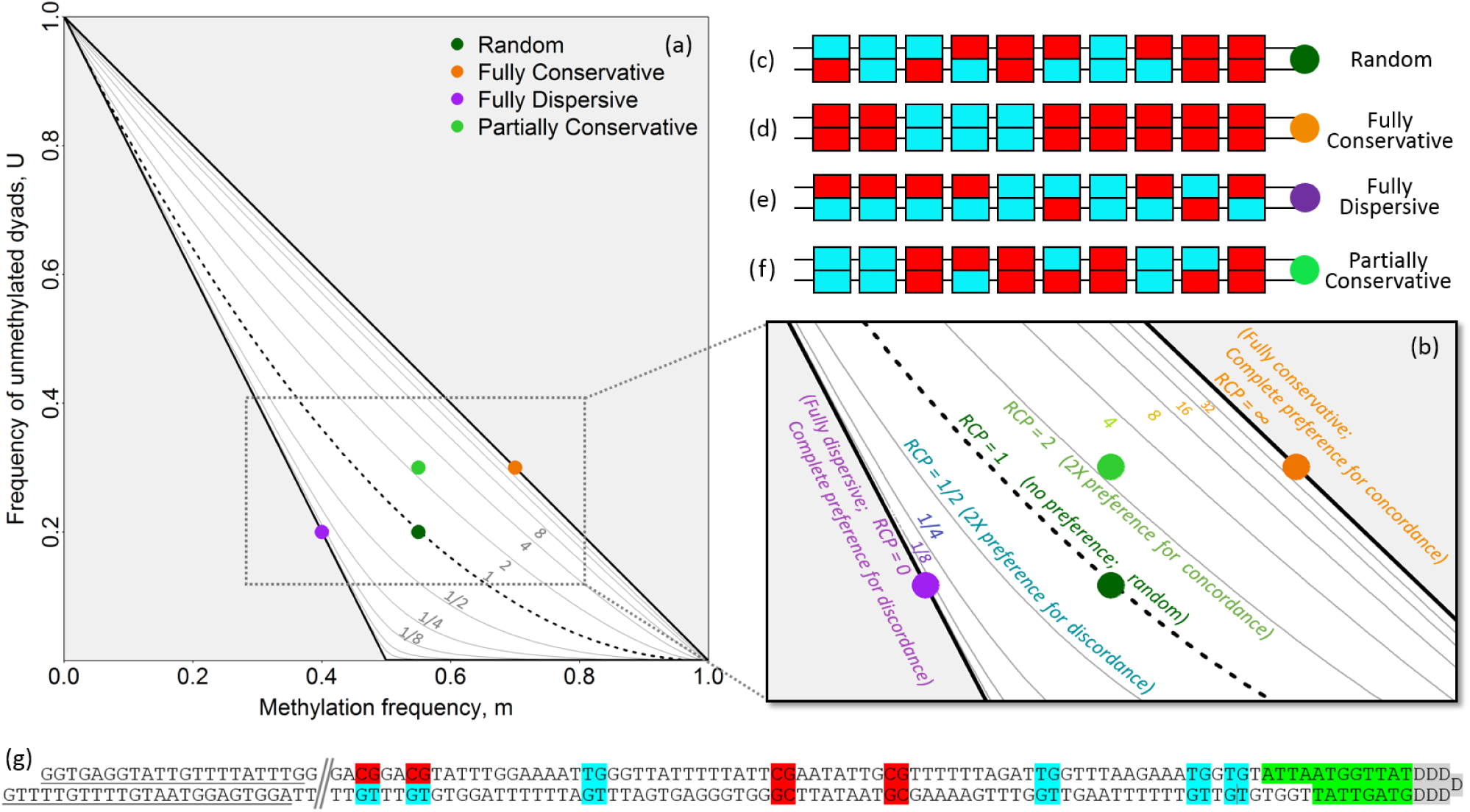
Characterizing Methylation Systems Using Double-Stranded DNA Methylation Patterns. **(a)** Frequencies of methylated cytosines (*m*) and unmethylated dyads (*U*) locate each data set on the continuum from complete preference for concordance to complete preference for discordance. Ratio of Concordance Preference (RCP) is indicated for each contour line. **(b)** Expanded view. For this schematic, individual double-stranded methylation patterns **(c-f)** are used to illustrate different methylation configurations that lie along this continuum. Individual patterns, with methylated and unmethylated cytosines indicated in red and blue, respectively, can reflect **(c)** a random methylation system; **(d)** a fully conservative system, with complete preference for generating concordant dyads; **(e)** a fully dispersive system, with complete preference for generating discordant dyads (partial preference for dispersive placement is also possible); or **(f)** a partially conservative system, with more concordant dyads than expected under random processes, but fewer than expected under fully conservative processes. **(g)** A representative partial double-stranded DNA methylation pattern collected using hairpin-bisulfite PCR. The experiment-specific batchstamp is shown in green, and can be used to monitor for PCR contamination; the molecule-specific barcode shown in gray, generalized as “DDDDDDD”, can be used to identify redundant sequences. The batchstamp and barcode are encoded by the hairpin oligonucleotide used to join the top and bottom strands. Primer-binding sites are underlined at the left end of the molecule.

One set of deviations from the random expectation is characterized by preference for concordant placement of methyl groups, such that the two classes of concordant dyads — fully methylated and fully unmethylated — are more frequent than expected under the random model. This situation occurs under conservative systems of methylation where strong contributions from maintenance-like processes, such as the activity of Dnmt1 in mammals [11, 13, 24], lead to high frequencies of concordant dyads. In the extreme form of this deviation from random, methyl groups are observed only in fully methylated dyads (Fig 1d), such that unmethylated dyads occur at frequency *U* = 1 *- m* (upper diagonal line in Fig 1a-b).

The other set of possible deviations from random is characterized by preference for discordant placement of methyl groups, leading to an overabundance of hemimethylated dyads. This situation occurs under dispersive systems of methylation such as those that yield transient hemimethylation following DNA replication and prior to daughter-strand methylation, and perhaps in genomic regions undergoing demethylation during periods of epigenetic transition. When methylation is maximally dispersive and methylation frequency *m* is less than 0.5, all dyads with methylation will be hemimethylated (Fig 1e), such that *U* = 1-2*m* (lower diagonal line in Fig 1a,b); when *m* is greater than 0.5, not all methyl groups can be accommodated in hemimethylated dyads, and so a combination of hemimethylated and fully methylated dyads — but no unmethylated dyads — is expected (lower horizontal line in Fig 1a).

The two extreme deviations from random form the boundaries of the comprehensive space of possible configurations of methylation at symmetric motifs (Fig 1a). Sets of double-stranded methylation patterns fall on the continuum between the extreme expectations (Fig 1b), and can be located within this space to characterize the strategy of information transfer employed to give rise to a given data set, ranging from conservative to dispersive.

As noted above, a system with an RCP value of 1 has no preference for either concordance or discordance of methylation, and is analogous to the distribution of genotype frequencies at a two-allele locus in a population that is at Hardy-Weinberg equilibrium [21, 22]. An RCP value of 2 indicates two-fold preference for concordance, while an RCP of 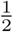 indicates two-fold preference for discordance. RCP approaches infinity for systems that have complete preference for concordant dyads. At the other extreme, RCP approaches 0 (i.e.,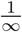) for systems that have complete preference for discordant dyads.

For the examples analyzed here, data for different loci and cells range from complete concordance to near-random, along the RCP spectrum. Complete discordance is found as a transient condition of adenine methylation at the *ori* locus in *Escherichia coli*, and serves to regulate the timing of reinitiation of DNA synthesis [25, 26]. Adenine methylation in *E. coli* generally occurs at symmetric sites, such as the GATC motif within the *ori* locus, and can be assessed by PacBio sequencing [27]. Thus, a broad spectrum of concordance preference can exist in organisms, and can be quantified and evaluated by RCP.

For large and intermediate-size data sets, the resolution of RCP is high across the range of possible methylation frequencies, although the resolution declines as *m* approaches 0 or 1, such that RCP cannot be inferred for completely methylated or unmethylated genomic regions. Nonetheless, RCP can usually be inferred with high confidence using data from only a few hundred dyads. Our new approach therefore requires far fewer sequences to estimate concordance preference than do methods that focus on inferring rates for specific enzyme activities [24, 28].

We apply RCP to investigate further the conclusion of Shipony *et al.* [16] that methylation in cultured stem cells is dominated by non-conservative processes, with little or no preference for concordance. Using double-stranded methylation patterns collected by our group, by Arand *et al.* [14, 17], and by Zhao *et al.* [15], we assess and compare methylation concordance in cultured human and murine stem cells, as well as in murine cells undergoing early developmental transitions that give rise to totipotent embryonic cells.

### Differentiated Cells Strongly Prefer Concordant DNA Methylation

Our previous work with human single-copy loci in uncultured, differentiated cells revealed a substantial role for maintenance methylation, a conservative process, with a comparatively minor role for non-conservative *de novo* processes [24, 28]. We therefore anticipated that RCP analysis of double-stranded methylation patterns from such cells would indicate substantial preference for concordant methylation states. Data published previously for *G6PD*, *FMR1*, and *LEP*, in uncultured differentiated cells and new data presented here for *FMR1* in cultured, human differentiated cells represent blood, connective, and adipose tissues. We applied RCP analysis to these data sets and found 13.2- to 85.7-fold preferences for concordant methylation. This confirms, as anticipated, that methylation is predominantly conservative in these differentiated cells (Fig 2a; see table accompanying Fig 2 for approximate 95 %CIs). We note that there is a good correspondence between RCP and hemi-preference ratio, a statistic we computed for the same data sets in the previous study [24] (further discussion in S2 Text).

**Figure 2.**
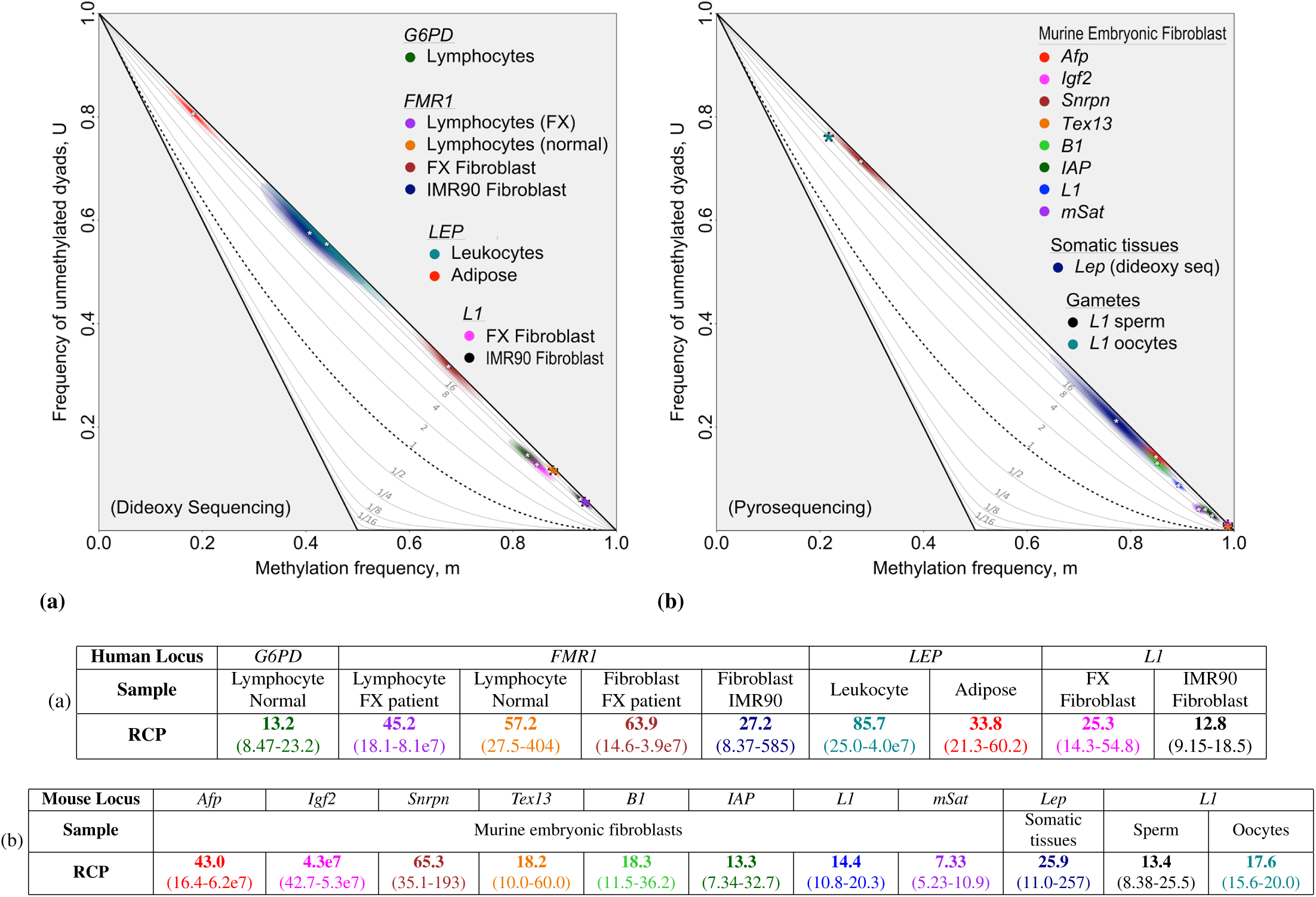
Inferring RCP for loci in differentiated human and murine cells. Methylation in human and murine differentiated cells was consistently inferred to have strong contributions from conservative processes, using data sets that span a wide range of methylation frequencies. For each locus or multi-copy family, we inferred the RCP point estimate of the *m*, *U* pair and the two-dimensional confidence region, determined by the uncertainty in the two variables (S7 Text). The intensity of coloration at a given point in a confidence region reflects the confidence level at that point. The *m*, *U* point estimates for most data sets are indicated with white asterisks, and the corresponding RCP values are given in the associated table. A larger, colored asterisk is used when the confidence interval of a data set is too small to be readily visible. RCP point estimates and bias-corrected bootstrap confidence intervals are shown in figure-associated tables. **(a)** Three single-copy human loci — *G6PD*, *FMR1*, and *LEP* — and one human repeat family, *L1*, all from various tissues as indicated. **(b)** Four single-copy loci and four repeat families – *Afp*, *Igf2*, *Snrpn*, *Tex13*, *B1*, *IAP*, *L1*, and *mSat* – from murine embryonic fibroblasts, one single-copy murine locus – *Lep* – from somatic tissue, and one repeat family – *L1* – from murine gametes. Data in (a) were collected using hairpin-bisulfite PCR and dideoxy sequencing, and taken from published [24, 28–30] and previously unpublished work (Table S3). Data in (b) were collected by Arand *et al.* [14] using hairpin-bisulfite PCR and pyrosequencing, with the exception of murine *Lep* for which data from somatic tissues were collected by Stoger [29], using dideoxy sequencing. Our analyses of these data sets applied bootstrapping approaches and accounted for inappropriate and failed conversion of methylcytosine using methods described in Supporting Information (S4 Text). Dyad counts, conversion-error rates, and inferences for methylation frequencies and RCPs are summarized in Tables S3, S4.

We also found a substantial role for conservative methylation processes at single-copy loci in both cultured and uncultured murine differentiated cells. Data sets from Arand *et al.* [14] for *Afp*, *Igf2*, *Snrpn*, and *Tex13* from murine embryonic fibroblasts (MEFs), and from Stöger [29] for *Lep* from somatic tissues, gave RCP point estimates indicating a greater than 18-fold preferences for concordant methylation (Fig 2b).

Do multi-copy sequence families also have high preference for concordant methylation in differentiated cells? We inferred RCP for four repeat families – *B1*, *IAP*, *L1*, and *mSat* – using murine data collected by Arand *et al.* [14]. Three of these families were found to have preference for concordant methylation in the same range inferred for single-copy loci (RCP point estimates between 14.4 and 19.3; Fig 2b). The fourth – *mSat* – had an RCP estimate of 7.33, lower than other families and single-copy loci examined in MEFs, but still indicative of strong preference for concordant methylation. For human cells, data from two independent lines of cultured embryonic fibroblasts were available for the repeat family *L1*. Inferred RCP values were within the range found for single-copy loci in both human and murine differentiated cells (Fig 2a).

Overall, we find appreciable preference for concordance across a diverse group of data sets from differentiated cells. These sets span a more than five-fold range in methylation frequency, underscoring the independence of RCP from *m*, and, more generally, highlighting the capacity of methylation systems to propagate specific epigenetic states, even when methylation is sparse. We conclude that preference for concordant methylation, albeit to variable degrees, is present in differentiated cells across broad classes of genomic elements, cell and tissue types, and culture states.

### Concordance Preference is Reduced but still Substantial in Cultured Stem Relative to Differentiated Cells

We next ask whether substantial preference for concordance, as we infer above for differentiated cells, is also evident in data from cultured stem cells. In doing so, we compare our findings using RCP to the expectation from Shipony *et al.* [16] that methylation in such stem cells occurs primarily through non-conservative, random processes.

The broadest data set available for our analysis comes from the near-genome-wide double-stranded methylation data presented by Zhao *et al.* [15]. These data give an inferred RCP of 5.22 for “all CpGs” in DNA from undifferentiated, cultured murine ES cells (Fig 3a; Table S5). For other classes of genomic elements in these near-genome-wide data [15], we infer RCP values of 4.31 or greater (Table S5). These RCP values are significantly higher than 1, the value predicted under Shipony *et al.*’s proposal of dynamic methylation (*p <* 10^-16^, maximum likelihood comparispon tests (MLCTs)).

**Figure 3.**
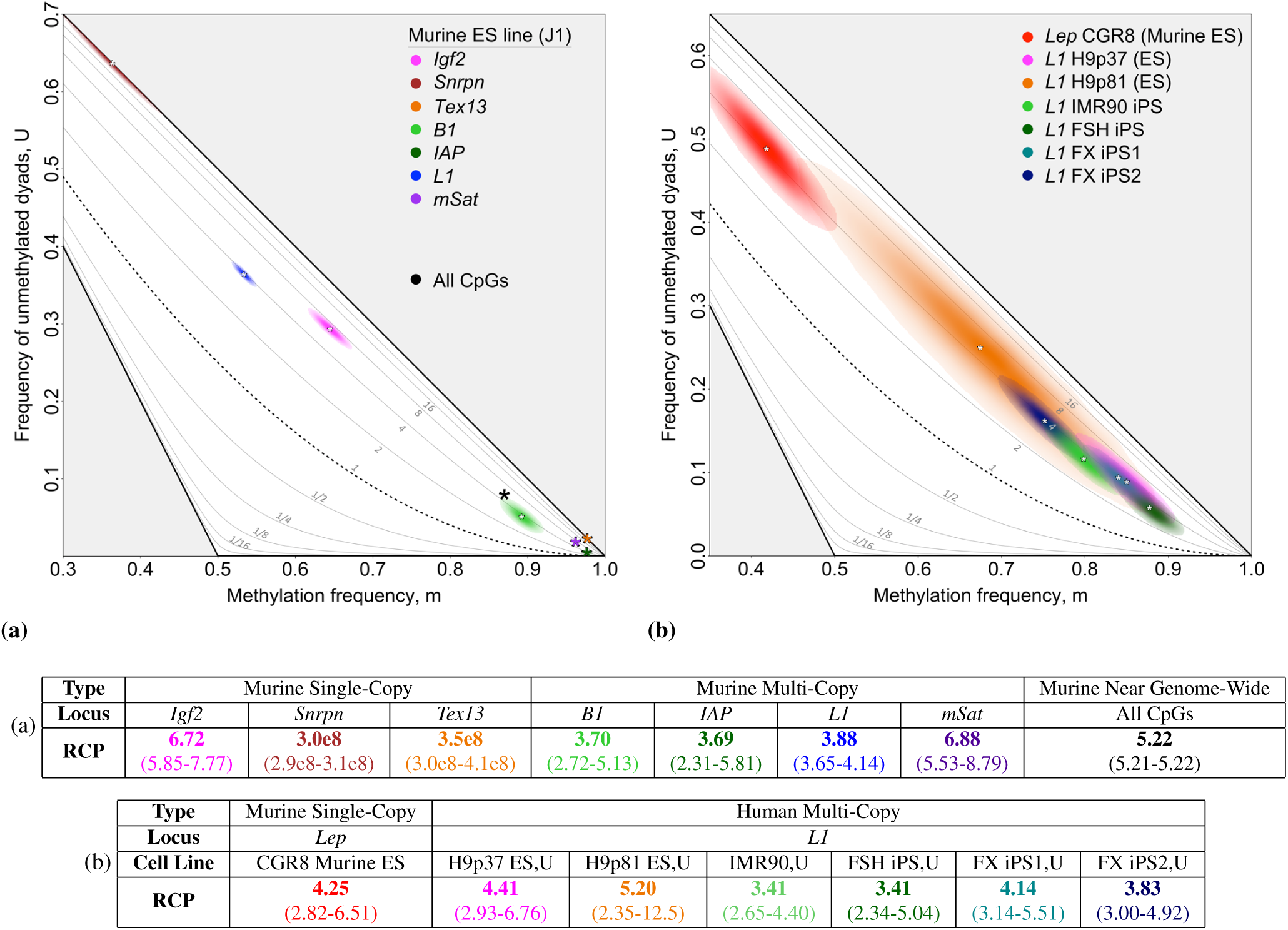
Inferring RCP in undifferentiated human and murine stem cells. Methylation patterns in undifferentiated, cultured human and murine stem cells were consistently inferred to have substantial contributions from conservative processes, with concordance greater than the random expectation. RCP point estimates and biased-corrected bootstrap confidence intervals are shown for individual loci and “All CpGs”. **(a)** Three single-copy loci – *Igf2*, *Snrpn*, and *Tex13* – as well as four multi-copy loci – *B1*, *IAP*, *L1*, and *mSat* – from murine ES J1 cells were assayed by Arand *et al.* [14]. “All CpGs” data, collected by Zhao *et al.* [15]), reflect methylation at 17.3% of CpG dyads in the murine genome (Table S5). Data from both Arand *et al.* [14] and Zhao *et al.* [15] were collected using hairpin-bisulfite PCR and pyrosequencing. **(b)** Human *L1* and *LEP* data were collected using hairpin-bisulfite PCR and dideoxy sequencing (data from published [29] and previously unpublished work (Table S3)). Our analyses of these data sets applied bootstrapping approaches and accounted for inappropriate and failed conversion of methylcytosine using methods described in Supporting Information (S4 Text). Dyad counts, conversion-error rates, and inferences for methylation frequencies are given in Tables S3, S4, and S5.

We next ask whether our inference of appreciable concordance preference in the murine ES cell line used by Zhao *et al.* [15] reflects a general property of cultured lines of undifferentiated stem cells, both murine and human. For the murine ES line, J1, double-stranded methylation data collected by Arand *et al.* [14] were available for four single-copy loci and four repeat families. Seven of the eight genomic regions – *Igf2*, *Snrpn*, *Tex13*, *B1*, *IAP*, *L1*, and *mSat* – had RCP values greater than 3.69 (with minimum 95%-CI lower bound of 2.31), still indicative of substantial preference for concordant methylation (Fig 3a). One single-copy locus, *Afp*, had a methylation level too high, 0.99, to permit reliable inference of RCP. Murine double-stranded methylation patterns for the four repeat families were available for two more stem cell lines, E14 and WT26 [14]. These additional repeat-family data sets, too, had RCP values significantly greater than 1, although one data set, that for *mSat* in WT26, had an RCP value closer to 1 than did others (*p* = 0.045, one-tailed BT). Data were available for a single-copy locus, *Lep*, for a fourth murine ES line, CGR8. Here, too, RCP was significantly greater than 1 (*p <* 10^-16^, one-tailed BT).

Human stem cell lines also followed the pattern of preference for concordant methylation. All six of the human stem and iPS cell lines that we examined, when grown under non-differentiating conditions, gave RCP point estimates for the repeat family *L1* that are between 3.41 and 5.20. For all of these cell lines, outer bounds of the approximate 95% confidence intervals fall between 2.34 and 12.53 (Fig 3b; Table S3). Together, these values reveal concordance preference that is reduced relative to differentiated cells, but still greatly exceeds expectations under random placement of methyl groups (p < 10^-16^, one-tailed BT).

We now consider the possibility that spontaneous differentiation had produced subpopulations of cultured stem cells that might account for the inference of RCP values substantially greater than 1 at the seven different loci and genomic elements examined. Our calculations revealed that a possible subpopulation of differentiated cells operating at much higher RCP than that of undifferentiated cells would need to comprise more than 50% of the population to account for our finding (S9 Text). Morphological inspection of the cultured human stem cells under non-differentiating conditions did not suggest the presence of a substantial subpopulation of differentiated cells in any of these lines.

We conclude that RCP values significantly greater than 1 are a consistent feature of cultured embryonic stem cells, and hold across a broad set of stem cell lines, genomic locations and element categories.

### Preference for Concordance is Minimal or Absent in ES Cells Deficient in Maintenance Methylation Activity

Our finding of substantial preference for methylation concordance in data from cultured, undifferentiated stem cells contrasts with the inference of Shipony *et al.* [16] that DNA methylation in such cells is dominated by non-conservative, random processes. This disparity led us to ask whether our approach here for data acquisition and analysis is indeed capable of identifying sets of methylation patterns established under exclusively random processes, which are expected to yield RCP values of 1 (see “Ratio of Concordance Preference is Defined …”, above).

To assess this capacity, we consider methylation patterns from two murine embryonic stem cell lines that have impaired maintenance methylation: a *Dnmt1* knockout (KO) line and an *Np95* KO line. The Dnmt1 enzyme is principally responsible for addition of methyl groups to daughter-strand CpGs complementary to CpGs methylated on the parent strand [11, 24, 32]; Np95 facilitates interaction of Dnmt1 with these hemimethylated sites [33]. Absence of either protein is therefore predicted to markedly diminish maintenance activity. If our approach is able to detect essentially random placement of methyl groups, RCP values in these knockout lines should be *∼*1 for loci for which Dnmt1, its actions facilitated by Np95, is principally responsible for conservative methylation.

Significant reductions in RCP were inferred for all single-copy loci and repeat families examined in *Dnmt1* and *Np95* KO lines [14, 31], compared to the parent stem-cell lines. Some reductions were sufficient to bring RCP values in the knockout lines to that expected for random placement of methyl groups: one single-copy locus – *Afp* – in the *Dnmt1* KO line and one repeat family – *B1* – in both knockout lines had RCP values not significantly different from 1 (*Afp* in *Dnmt1* KO: 1.16, *p* = 0.10; *B1* in *Dnmt1* KO: 1.14, *p* = 0.17; *B1* in *Np95* KO: 1.02, *p* = 0.38; one-tailed BTs; Fig 4). These findings in the two mutant cell lines thus reveal that RCP analysis is, indeed, able to detect methylation established with random placement of methyl groups, and thus with little or no preference for concordance or discordance.

**Figure 4.**
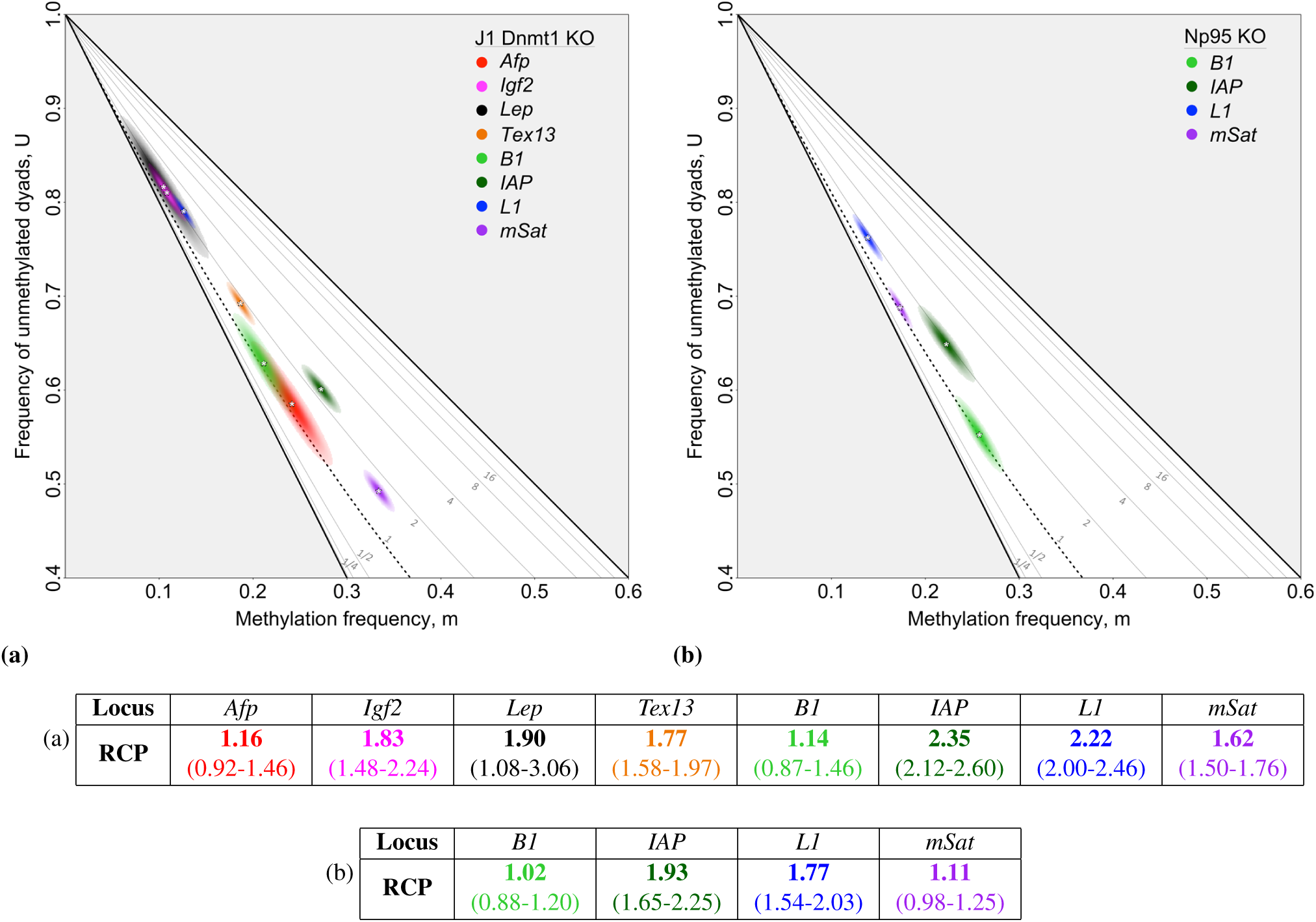
Inferring RCP in *Dnmt1* knockout and *Np95* knockout ES cells. (**a**) In the absence of Dnmt1, the primary maintenance methyltransferase, RCPs inferred for cultured murine ES cells were close to 1, the random expectation, at some loci. Data from *Lep*, collected using hairpin-bisulfite PCR and dideoxy sequencing, are from Al-Alzzawi *et al.* [31]. Data from seven additional loci are from Arand *et al.* [14], who used hairpin-bisulfite PCR and pyrosequencing. Our analysis of these published data revealed two of the eight loci analyzed — *Afp* and *B1* — to have very low RCP values not significantly different from 1. (**b**) In the absence of Np95, a protein critical for recruiting Dnmt1 to hemimethylated regions in newly replicated DNA, one of the four loci analyzed — *B1* — was inferred to have a very low RCP value, not significantly different from 1. Our analyses of these data sets from Arand *et al.* [31] applied bootstrapping approaches and accounted for inappropriate and failed conversion of methylcytosines using methods described in S4 Text. Point estimates and approximate 95% confidence intervals on RCP are given above, and also along with conversion-error-rate estimates in Tables S3 and S4.

The ability of RCP to detect random methylation has important implications for our work. First, we can conclude that our inference of persistent preference for concordant methylation in cultured stem cells reflects a *bona fide* property of those cells, rather than an artifact of our approach. Second, we can infer from our finding of RCP *>* 1 for nine of the twelve data sets examined in the *Dnmt1* and *Np95* KO lines that methyltransferases other than Dnmt1 can contribute to conservative methylation. This inference is consistent with earlier conclusions that contributions of Dnmt3s can include low levels of maintenance activity [19, 24, 34].

### Concordance Preference Increases upon Differentiation of ES Cells, and Decreases upon Dedifferentiation

Our initial examination of RCP values in differentiated cells as compared to cultured stem cells suggests that RCP is altered through the differentiation process (Fig 2 and Fig 3). Would significant RCP increases be observed for individual cell lines transitioning between differentiation states? We first asked whether RCP values change when undifferentiated human ES and iPS cells are grown under differentiating conditions (see Materials and Methods). We inferred RCP at the promoter of *L1* elements of cultured human iPS and ES cells, inferring values for two different passages of the latter cell line. Upon differentiation, RCP values for all three of these cell lines increased significantly (*p* = 0.004, H9p37; *p* = 0.025, H9p81; *p* = 0.0005, FSH iPS; two-tailed PTs), and approached the lower boundary of the confidence region inferred for single-copy loci in differentiated somatic cells (Fig 2 and Fig 5a). Using near-genome-wide data for cultured murine cells [15], we inferred significant RCP increases upon cell differentiation for most genomic elements (*p <* 10^-16^, MLCTs), with the exception of low-complexity and satellite DNAs (Table S5). These RCP increases were greatest at promoters, CG islands, and CG shores, and are more modest at other regions. We conclude that the onset of differentiation in cultured human and murine cells is associated with a shift towards a greater role for conservative processes.

**Figure 5.**
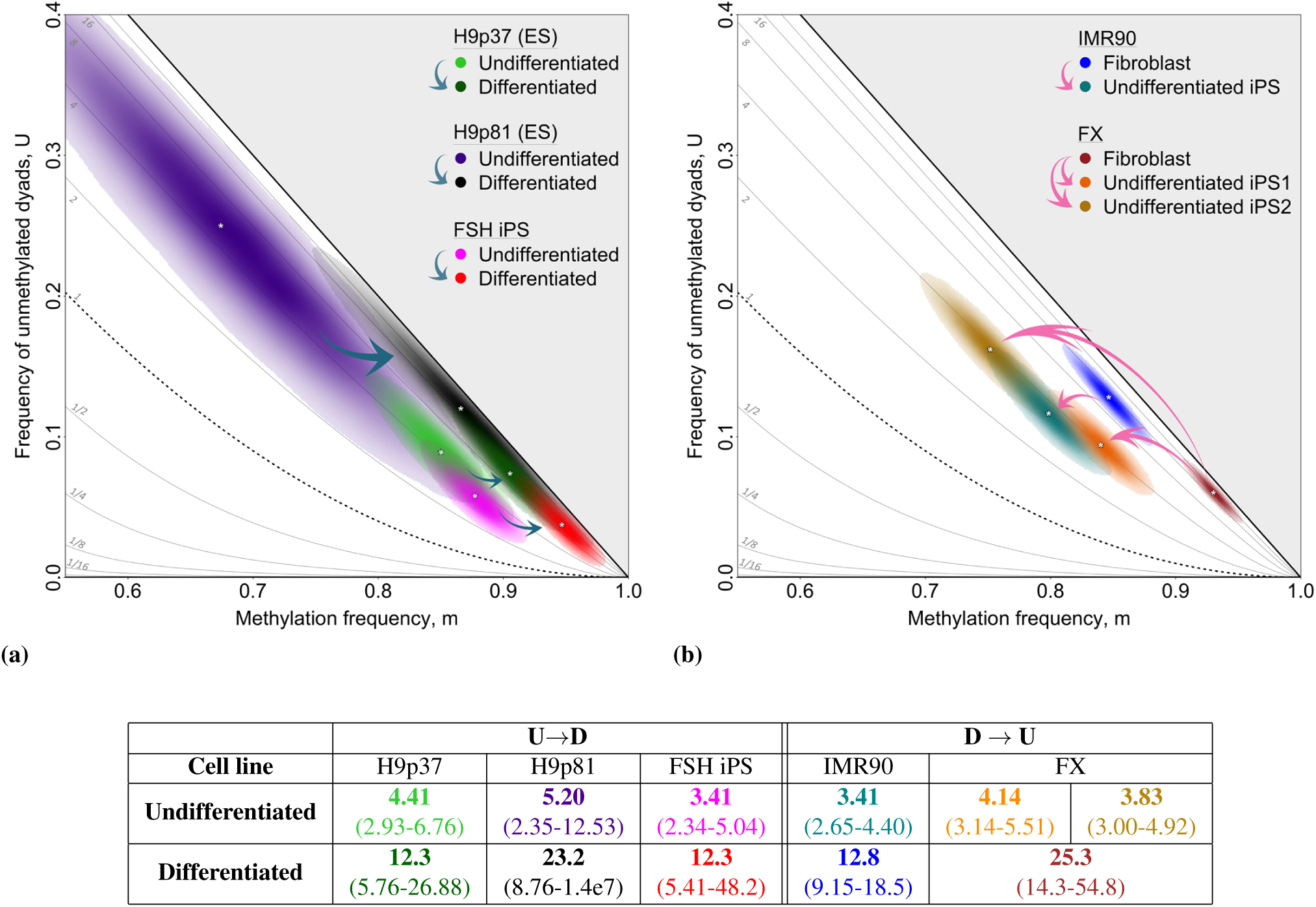
Shifts in RCP of *L1* elements upon differentiation of cultured human ES cells and dedifferentiation of cultured fibroblasts. RCP values of *L1* elements in cultured stem cells grown under non-differentiating conditions are compared to those for the same cells grown under differentiating conditions. RCP values for progenitor fibroblast lines are compared with those for their descendent iPS cells. Blue arrows indicate differentiation and pink arrows indicate dedifferentiation. **(a)** Undifferentiated cells from each of three human ES and iPS lines had only moderate preference for concordance. Upon differentiation, their RCP values shifted in parallel toward stronger preference for concordance. **(b)** Differentiated human fibroblast lines had substantial preference for concordance. Upon dedifferentiation to iPS cells, RCP values were reduced, indicating diminished preference for concordance. We present data for two different iPS lines established independently from the fibroblast line, FX. These iPS lines differed in their methylation frequency, *m*, at *L1* elements, but had similar RCP values. Some data were previously shown in Fig 2 and Fig 3, and are included again here to illustrate more directly the relationships of *L1* methylation in the differentiated and undifferentiated cultured cells. We applied bootstrapping approaches and accounted for inappropriate and failed conversion of methylcytosine using methods described in S4 Text. Point estimates of RCP, with approximate 95% confidence intervals, are shown above, and also with dyad counts, conversion-error rates, and inferences for methylation frequencies in Tables S3 and S4.

Does the dedifferentiation that occurs in culture upon production of an iPS line from a differentiated cell have an opposite effect on concordance preference? To address this question, we compare methylation at *L1* elements in three iPS lines to that in the two cultured human fibroblast lines from which they were derived. As predicted, RCP values for all three iPS lines were much reduced compared with values observed for the parent fibroblast lines (*p <* 10^-16^, two-tailed PTs; Fig 5b). Dedifferentiation in tissue culture is thus associated with a shift in DNA methylation toward a greater role for non-conservative processes. It will be useful to investigate whether changes in methylation systems as measured by RCP drive or merely reflect the cellular differentiation process.

### Murine Embryos Mirror the Transitions in Concordance Preference Observed in Cultured Cells

The *∼*3-fold-or-greater preference for concordant methylation we infer in many different cultured ES and iPS cell lines (Fig 3a; Table S3) far exceeds the concordance expected under the null hypothesis that methyl groups are placed at random. Here we ask whether this appreciable preference for concordance is an artifact of growing stem cells in culture, or whether it is shared by uncultured stem cells taken directly from an embryo.

We first consider whether totipotent cells from an embryo have evidence of conservative processes. We applied RCP to double-stranded methylation patterns collected by Arand *et al.* [17] for three multi-copy loci in mouse embryos: *L1*, *mSat*, and *IAP*. Our analyses revealed that these totipotent embryonic cells, from post-replicative zygote to morula stage (through 3 days post conception, dpc), also exhibit moderate preference for concordance. Each of the eighteen data sets we considered yielded RCP point estimates greater than 1, and confidence intervals that exclude 1 (*p <* 0.005; Fig 6a; Table S4).

**Figure 6.**
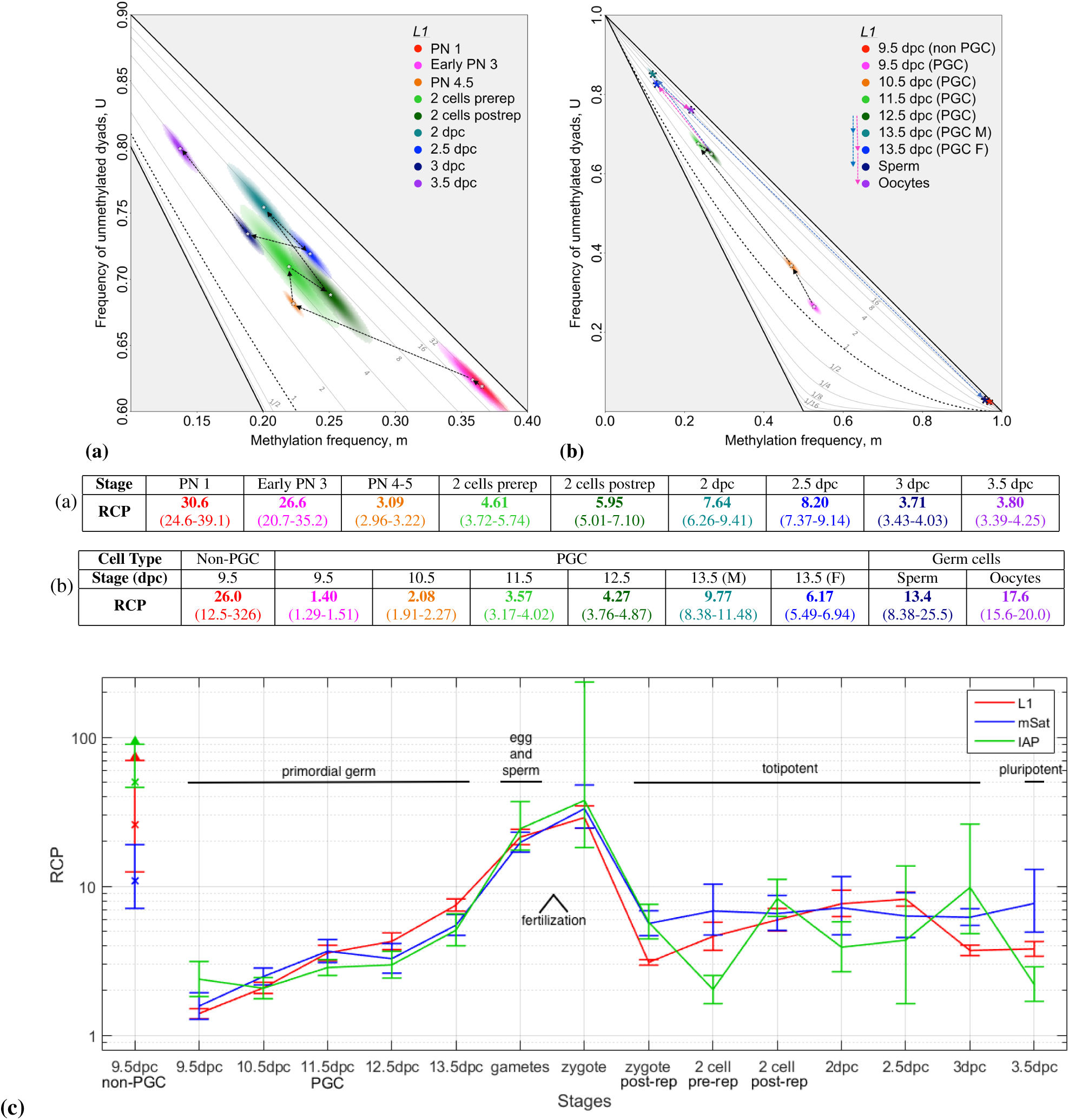
Shifts in RCP during murine embryonic and germ-cell development. Major transitions in RCP values occur during early embryonic and primordial germ cell (PGC) development. RCP point estimates and approximate 95% bias-corrected bootstrap confidence intervals are given in the associated tables and in (c). **(a)** Transitions from early pronuclear stages (1 and 3) to late stages (4-5) were accompanied by a sharp decrease in RCP at *L1* elements, to a level similar to that observed in cultured stem cells (Fig 3). Further embryonic development was accompanied by minor increases and subsequent decreases in methylation and RCP values. **(b)** In PGCs at the earliest stage for which data are available, 9.5 days post conception (dpc), *L1* elements had RCP values that were unusually low but still significantly greater than 1 (*p <* 10^-16^). RCP values increased during PGC maturation to stage 13.5 dpc even as methylation frequencies decreased by more than 50%. RCP values for eggs and sperm, shown previously in Fig 3, are included here to highlight the transition from early primordial germ cells to terminally differentiated gametes. **(c)** Tracking RCP at repeat families during development. We inferred RCP for the embryonic and differentiated stages for which data were published by Arand *et al.* [17]. The topology of our longitudinal RCP plot highlights the transitions also evident in Arand *et al.*’s plot of the percentage of hemimethylated CpG dyads relative to all methylated CpG dyads (Figure 6b in [17]). Their metric captures relative shifts in concordance, but, in contrast to RCP, does not include a null model enabling quantitative comparison of inferred concordance across data sets with disparate methylation frequencies [17]. Point estimates and approximate 95% confidence intervals for all but one data set were estimated by bootstrapping and applying the BCa correction (S7 Text). Dyad counts, methylation frequencies, and conversion error rates are in Table S3. In cases where data were available for multiple replicates, dyad frequencies were pooled. For one data set, 3 dpc at *mSat*, double-stranded sequences and methylation patterns were not available to us for bootstrap analysis; we therefore used a likelihood-based method for the estimation of confidence intervals (S8 Text).

Pluripotent stem cells from mouse embryos (gastrula, 3.5 dpc) also exhibit moderate preference for concordant methylation for all three of the multi-copy loci examined (*p <* 10^-16^; Fig 6a; Table S4). Thus, we conclude that moderate preference for concordance is an epigenetic feature of uncultured embryonic stem cells of disparate developmental potential, and is not an artifact of the establishment of embryonic stem cells in culture.

Though RCP values for stem cells from embryos clearly indicate some preference for concordance, the extent of this preference is lower than for differentiated cells at most of the loci examined (Fig 2 and Fig 5). Does this lower preference for concordance originate in gametes, or does it instead arise post-fertilization? To address this question, we expanded our inference of RCP values for sperm and oocytes from just *L1* (Fig 3) to all three loci analyzed by Arand *et al.* [17]. At the three multi-copy loci, RCP values for gametes are within the range observed for other differentiated cells, with average point estimates ranging from 13.4 to 45.0 (Fig 6b; Table S4). This high preference for concordance in gametes implies that the lower RCP values characteristic of zygotes and stem cells must arise post-fertilization rather than in gametes.

To pinpoint the timing of this transition to lower RCP values observed in stem cells, we consider data from post-fertilization nuclei and cells [17]. Data available for pronuclear stages 1, 2, and 3 revealed high RCP values, similar to those observed in gametes. In pronuclear stages 4-5, however, there was an abrupt transition to lower RCP values in the range observed for totipotent stem cells (Fig 6; Table S4).

Is this transition dependent on the DNA replication event that occurs from pronuclear stage 3 to stages 4-5? To address this question, we assess data from Arand *et al.* [17], in which aphidicolin was used to block DNA replication in the fertilized egg. Our RCP analysis reveals that methylation patterns at *L1* and *mSat* in these treated cells, while having somewhat lower RCP values relative to pronuclei at earlier stages, had not undergone the major reduction in RCP that we infer for unmanipulated pronuclei at stages 4-5 (*p <* 10^-16^, two-tailed PTs; Table S4). Thus, the shift to lower RCP in stem cells following fertilization appears to require either DNA replication or some later event that is itself replication-dependent. This conclusion is consistent with the inference of Arand *et al.*, using a different metric [17], that DNA replication in the zygote plays a pivotal role in methylation dynamics.

This change from high RCP values in gametes and early pronuclei to lower values in post-replicative zygotes and descendant stem cells was markedly abrupt. Are other major transitions in RCP similarly abrupt, or do some occur gradually, perhaps over many cell divisions? Murine primordial germ cells (PGCs), in their maturation to differentiated gametes, offer an opportunity to approach this question [35, 36]. Using data from Arand *et al.* [17], we infer that RCP values for PGCs increased by factors ranging from 2.5 to 5 during their progression from 9.5 dpc to 13.5 dpc. This increase was not sudden, but occurred over the four-day period, and so spanned an interval of substantial cell proliferation [35] (Fig 6b). The murine embryo data thus provide examples of both abrupt and gradual transitions in RCP through development (Fig 6c).

The RCP values for PGCs at 9.5 dpc, the earliest stage for which data were reported by Arand *et al.* [17], were strikingly low: 1.40 for *L1*, 1.57 for *mSat*, and 2.38 for *IAP*. Nonetheless, confidence intervals for all three loci indicated RCP values greater than 1 (*p <* 2×10^-5^), the value expected expected under wholly random placement of methyl groups, indicating persistent, low-level preference for concordance. This low, residual preference for concordance in maturing PGCs perhaps reflects both the epigenetic memory needed to maintain the poised state of stem cells, and the epigenetic flexibility required for the production of differentiated gametes.

The first few rounds of PGC division involve developmental reprogramming and commitment [36], and establishment of lineage-specific gene expression patterns. The 3.5-fold average increase of RCP across these divisions (Fig 6; Table S4) mirrors the 3.6-fold average increase in RCP values that occur when cultured ES cells are subjected to differentiating conditions (Fig 5; Table S3).

There is, however, a critical difference between the trajectories for proliferating PGCs and differentiating ES and iPS cells in culture. When cultured cells were differentiated, their methylation frequencies increased. By contrast, methylation frequencies for PGCs declined across early rounds of division [37]. Seisenberger *et al.* [7], *Arand et al.* [17], and von Meyenn *et al.* [18] concluded that this reduction in methylation frequency is driven by partial impairment of maintenance methylation.

Our RCP framework permits a closer look at the likely extent of this proposed maintenance impairment. If maintenance methylation were completely absent and no other methylation processes were active, passive, fully dispersive demethylation would occur. This would halve methylation frequencies and leave methyl groups only in hemimethylated dyads, yielding an RCP value of 0. By contrast, data from PGCs yielded RCP values significantly greater than 0, and even 1. Indeed, only cells treated with S-Adenosylmethionine-ase (RCP point estimate: 0.20, approximate 95% confidence interval: 0.15 - 0.26; Table S4) yielded values close to 0. This is not surprising, as S-Adenosylmethionine-ase either impairs or eliminates a cell’s ability to methylate DNA, and so reveals RCP trajectories that would be observed with complete or nearly complete suspension of all methylation processes. Together these findings confirm that, while the RCP framework can detect very low RCP values, maturing PGCs retain conservative methylation processes, and that these processes occur at levels sufficient to outweigh any dispersive effects of passive demethylation.

## Concluding Remarks

Because RCP makes no explicit enzymatic or mechanistic assumptions about the methylation machinery, it permits quantification and comparison of strategies for symmetric methylation across cell types, developmental periods, and organisms, despite potential and likely differences in exact mechanisms. Application of RCP to double-stranded DNA methylation patterns reveals that preference for concordance in DNA methylation is a persistent though quantitatively variable feature of mammalian cells of disparate developmental potential. Specifically, we find that: (i) in cultured human and murine ES and iPS cells, preference for concordance is lower than in differentiated cells, but not absent; (ii) for cultured human stem cells, cellular differentiation is characterized by increasing preference for concordance, whereas, for cultured differentiated cells, dedifferentiation is characterized by declining preference for concordance; and (iii) during early murine development, transitions in RCP mirror those found in cultured cells, with pluripotent and totipotent stem cells showing appreciable concordance preference throughout. We also observe that substantial changes in RCP can be either abrupt, requiring only one DNA replication event, or gradual, occurring over multiple rounds of replication.

Although preference for concordance is substantial throughout early murine development, there is an instance of concordance preference near the expectation under entirely random processes. We infer RCP values close to, albeit significantly different from, 1 in the early primordial germ cell stage at the three repetitive element families examined by Arand *et al.* [17] (Fig 6). The instance of low, yet present, concordance preference may reflect both the epigenetic stability required to maintain the poised state of the stem cells and the epigenetic flexibility needed *en route* to production of functional gametes. Flexibility, indicated by RCP values near 1, may result from near-random processes or instead from a balance of conservative and dispersive methylation. Existing data and conclusions of Seisenberger *et al.* [7], *Arand et al.* [17] and von Meyenn *et al.* [18] are more consistent with the latter interpretation.

Our finding of moderate contributions from conservative DNA methylation processes in human and murine stem cells is seemingly contrary to the conclusion of Shipony *et al.* [16] that “dynamic” processes are dominant in cultured stem cells, even in regions where dense methylation is maintained. This apparent disparity may have arisen from differences between the temporal scales assayed by Shipony *et al.*’s approach and our own. The method of single-cell isolation and clonal expansion used by Shipony *et al.* estimates epigenetic memory from single-stranded data collected after 15 to 21 rounds of cell division. In contrast, our approach utilizes double-stranded DNA data to examine epigenetic memory over a single round of DNA replication. Evidence of preference for concordance, apparent in our comparison of DNA strands separated by one replication event, will be muted in comparisons of more distantly related molecules.

Short-term epigenetic memory, perhaps important for guiding cell-fate trajectories at early developmental stages, is at least partially achieved through preference for concordant DNA methylation. By contrast, over larger numbers of cell divisions, as sampled by Shipony *et al.* [16], such as for stem cells dividing in culture, preference for concordant methylation may be less important than other mechanisms of epigenetic memory. For example, regulation of promoter activity by DNA methylation can occur via an ensemble effect rather than by methylation of specific CG dyads within a promoter [38]. In such cases, propagation of exact methylation patterns may be less important than the density of methylation that influences gain or loss of methylation and states of transcriptional activity over many cell divisions [39]. Epigenetic mechanisms other than DNA methylation also contribute to epigenetic memory at various timescales. RCP analysis in combination with histone-modification data from ENCODE [40] and Roadmap [41] will provide unprecedented opportunities to infer interactions between DNA-methylation machinery and histone modification, the developmental timing of epigenetic stability, and its variation across the genome.

The value of RCP analysis will be enhanced and broadened by emerging DNA sequencing technologies that yield longer, more informative double-stranded methylation patterns. Longer sequence reads will enable inference of RCP for single cells, permitting study of cell-cell epimosaicism, such as arises in cancer [2] and other syndromes characterized by epigenetic heterogeneity and change [30, 42]. Some of these methods also reduce data corruption arising through errors in bisulfite conversion and amplification, and can distinguish between methyl- and hydroxy-methyl cytosine [43, 44]. High-resolution RCP estimates available through these advances will provide new insight into the flexibility and potential sensitivity of individual loci and cell types to environmental conditions encountered during embryogenesis and beyond.

## Materials and Methods

### Mathematical Foundations of the RCP framework

Overview of the mathematical foundation is given in Results and Discussion, and developed further in S1 Text.

### Human Cells and Culture Conditions

The six human ES and iPS cell lines for which we collected methylation patterns were derived from either embryos or fibroblasts described as normal [45] or from fibroblasts of individuals with disorders not known to affect the basic biochemistry of DNA methylation.

Many of the sets of human DNA methylation patterns analyzed here were presented in previous publications, which include information on University of Washington Human Subjects approval for collection and use. These data include *G6PD* and *FMR1* from leukocytes of normal individuals [24, 28]; *FMR1* from males with fragile X syndrome [30]; *LEP* from male leukocytes and from female lymphocytes and adipose tissue [24, 29].

Human methylation patterns presented here for the first time were collected from: (i) FX iPS cell lines 1 and 2, which were developed at the University of Washington ISCRM facility from fibroblasts (line GM07730, Coriell Cell Repositories, Camden, NJ) of a male with a fragile X “full mutation”, using published methods [46] ; (ii) iPS cell line IMR90, which was developed at the University of Washington ISCRM facility from the IMR90 somatic line established from fibroblasts (obtained from ATCC) of a normal female, using published methods [46]; (iii) FSH iPS cell line, which was developed at the University of Washington ISCRM facility from fibroblasts of an individual with Facioscapulohumeral dystrophy (FSHD), using published methods [47]; and (iv) H9 human ES cells from NIH Embryonic Stem Cell Registry (WA09, H9 number 0062).

Cells were cultured in Dulbecco’s modified Eagle’s medium/Ham’s F-12 medium containing GlutaMax supplemented with 20 percent serum replacer (SR), 1 mM sodium pyruvate, 0.1 mM nonessential amino acids, 50 U/ml penicillin, 50 *µ*g/ml streptomycin and 10 ng/ml basic fibroblast growth factor (all from Invitrogen), and 0.1 mM *β*-mercaptoethanol (Sigma-Aldrich). hESCs were grown on *γ*-irradiated primary mouse embryonic fibroblasts (MEFs) and passaged using dispase (1.2 U/ml; Invitrogen). They were passaged onto Matrigel (Corning) without feeders in mTeSR1 (Stem Cell Technologies) for the final passages prior to analysis. Cells were differentiated by passage onto Matrigel in Dulbecco’s modified Eagle’s Medium supplemented with 20 percent fetal bovine serum and pen/strep. Images of our cultured stem cells grown under differentiating conditions confirmed their pluripotency.

### Murine Cells and Culture Conditions

Methylation patterns from murine ES cells, and the origin and culturing of these cells, have previously been described [14, 15, 17, 31].

### Collection of Double-Stranded DNA methylation patterns using hairpin-bisulfite PCR

The DNA methylation patterns collected in our lab and analyzed here, both those published previously and those presented here for the first time, were collected using the hairpin-bisulfite PCR approach [13], with barcodes and batchstamps to authenticate each sequence [48]. Details for collection of each data set are given in Table S2.

The data presented by Arand *et al.* [14, 17], and Zhao *et al.*, [15] and analyzed here, were collected in the absence of molecular batch-stamps and barcodes, raising the possibilty that the reliably of those data sets is undermined by PCR clonality. However, both groups used alternate strategies that revealed that PCR clonality was not rampant. Zhao *et al.* found that essential features of data sets did not differ appreciably between the “real” data set collected conventionally, using PCR, and a test, PCR-free data set that excluded opportunities for clonality by including only one read from each locus, providing no evidence of impacts from PCR clonality (Hehuang Xie, personal communication). In turn, Arand *et al.* used molecular codes for several of their data sets, and, for the few data sets collected in the absence of such code, observed appreciable heterogeneity among patterns, also hinting that data were not appreciably impacted by clonality (Julia Arand, personal communication).

## Acknowledgments

We thank: Amos Tanay for suggesting a resolution between our results and those of Shipony *et al.*; three anonymous reviewers for insightful and constructive comments; Hehuang Xie, Ming-An Sun, and Julia Arand for generously sharing raw data from their published studies of methylation dynamics; Jeremy Johnson provided suggestions key to our data analysis; Stanley Gartler, Barbara Wakimoto, Matthew Stephens, Elizabeth Thompson, Carl Bergstrom, Frank Stahl, Jette Foss, Takuo Yamaki, Jesse McClure, Audrey Mat, Hyun Ji Noh, Steve Henikoff, Angela Liang, Alice Tao, Brooks Miner, Simina Popa, Rachana Kumar, and M. Goe for insightful and helpful comments.

## Supporting Information

**Figure S1. Markov chain used for the derivation of RCP.**

**Figure S2. RCP in *Dnmt3* knockout cells. (a)** RCP values were inferred at the *Lep* locus for wildtype, *Dnmt3a* KO, and *Dnmt3b* KO murine ES cell lines, using data from Al-Azzawi *et al.* [31]. **(b-i)** RCP values were inferred for eight additional loci for wildtype, *Dnmt3a* KO, *Dnmt3b* KO, and *Dnmt3a/b* double-KO murine ES cell lines, using data from Arand *et al.* [14].

**Table S1. Comparison of RCP values inferred here to the DNMT1 hemi-preference ratios (HPR) inferred by Fu *et al.* [24].** Point estimates and approximate 95% confidence intervals are shown for both statistics. Fu *et al.* provides only lower bounds of approximate 95% confidence intervals.

**Table S2. Hairpin-linkage and PCR conditions for collection of double-stranded DNA methylation patterns.** Entries shown are for patterns published here for the first time. R.E. refers to the restriction enzyme used to create the genomic overhang prior to ligation with a hairpin linker.

**Table S3. RCP values and associated approximate confidence intervals inferred for the 24 data sets from our labs.** We collected double-stranded DNA methylation patterns from two species — mouse and human — and several loci using bisulfite conversion under either low-molarity/temperature (“LowMT”) or high-molarity/temperature (“HighMT”) conditions [49]. For each data set, we counted methylated (*M*), hemimethylated (*H*), and unmethylated (*U*) dyads, and used these values to infer methylation frequency, *m*, unmethylated dyad frequency, *U*, and the ratio of concordance preference, RCP. Inferences for (*m, U*) and for RCP incorporated correction for conversion error, as described in S3 Text. We estimated approximate 95% confidence intervals by bootstrapping and applied the BCa correction to obtain bias-corrected confidence intervals and point estimates, as described in S7 Text. Both uncorrected and BCa-corrected intervals and point estimates are listed here.

**Table S4. RCP values and associated approximate confidence intervals inferred for data reported in Arand *et al.* [14, 17].** Dr. Julia Arand kindly shared raw double-stranded sequences for samples described in these publications. We accounted for failed and inappropriate conversion, as described in S3 Text, in our point estimates of *m*, *U*, and RCP. The inappropriate conversion rate of 0.031 was inferred because the bisulfite conversion conditions used by Arand et al. resembled lowMT conditions [49]. We estimated approximate 95% confidence intervals by bootstrapping and applied the BCa correction to obtain bias-corrected confidence intervals and point estimates, as described in S7 Text. Both uncorrected and BCa-corrected intervals and point estimates are listed here. For one sample, Arand *et al.* (2015) *mSat* 3dpc, only dyad counts, but no raw sequences, were available. We used the dyad counts to estimate the point estimate and the confidence interval for this sample while assuming independent sampling of dyads (S8 Text).

**Table S5. RCP values and associated approximate confidence intervals inferred for data reported in Zhao *et al.* (2014) [15].** Dr. Hehuang Xie kindly shared raw data on *M*, *H*, and *U* values for samples described in Figure S6 of Zhao *et al.*, and also provided information on error rates from bisulfite conversion: for the “Day 0” sample (cultured stem cells grown under non-differentiating conditions), the failed conversion rate was 0.011, and the inappropriate conversion rate was 0.0109; for the “Day 6” sample (cultured cells grown for 6 days under differentiating conditions), the corresponding error rates were 0.012 and 0.0099. Dr. Xie also commented, “‘All’ refers to the total CG dyads and ‘DNA’ refers to the CG dyads within ‘DNA repeat elements (DNA)’ annotated in the UCSC genome database.” For these data sets, whose individual sequence reads only contain 2 to 3 dyads on average, we assumed independent sampling of dyads to obtain the confidence intervals (S8 Text).

**Table S6. Pairwise comparisons of RCP values inferred for replicate samples of cultured human ES and iPS cells, and for mouse embryonic cells sampled at various developmental stages.** We applied our test for heterogeneity, as described in S5 Text, to assess evidence of significant differences between RCP values inferred from various sample replicates presented by Arand et al. (2015) [17]. A color gradient encodes approximate levels of significance, with dark green indicating non-significant differences, and red indicating highly significant differences.

### S1 Text: Deriving the Ratio of Concordance Preference, RCP

The goal of RCP is to quantify the extent to which the DNA methylation machineries that gave rise to each data set deviate from random expectations under the binomial distribution, as indicated by an over- or under-abundance of concordant dyads. Here we approach this problem by modeling the formation of concordant and discordant dyads as transitions between unmethylated and hemimethylated dyads and between hemimethylated and fully methylated dyads, without regard to the molecular processes that facilitate these transitions. We then take a mathematical approach to derive an expression for RCP.

We seek the equilibrium frequencies of fully methylated (*M*), hemimethylated (*H*), and unmethylated (*U*), dyads. Consider a continuous-time Markov chain operating on the probability distribution of the dyads *(M, H, U)*:

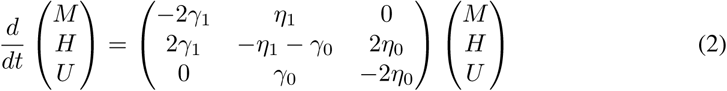

where *η*’s and *γ*’s represent the rates of methylation addition and removal, respectively, as shown in Fig S1.

We define RCP as the ratio between *β*_*c*_, the rate of dyad transitions yielding concordant dyads, and *β*_*d*_, the rate of dyad transitions yielding discordant dyads. We thus define 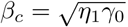, the geometric mean of the methyl-addition and methyl-removal rates yielding concordant dyads. Likewise, we define 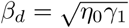.

We can then solve for the steady state distribution for the Markov chain in Equation (2) to arrive at the ratio. We can also express it in terms of *m* and *U*, as shown below.

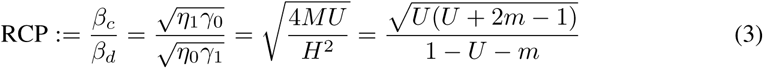

This formulation of RCP does not require the assumption that methylation frequency is constant over time. Here, “steady state” refers to the dyad frequencies expected under a given system of methylation processes, regardless of whether the methylation frequency is constant over rounds of cell division.

It is notable that RCP^2^ is 4*M U/H*^2^, which is expected to equal 1 under the Hardy-Weinberg equilibrium [21], if dyad frequencies are considered as genotype frequencies of a gene with two alleles. Following this, RCP can be considered as a measure of deviation from the null equilibrium.

RCP is therefore a metric for the degree to which the system of methylation processes prefers concordance (RCP *>* 1), discordance (RCP *<* 1), or, possibly, exhibits no preference in either direction (RCP = 1). If we set RCP = 1 and solve this expression for *U*, we find that *U* = (1*-m*)^2^. This is consistent with the expectation under the binomial distribution that RCP will be 1 when there is no preference for either concordance or discordance. As RCP approaches *∞*, *U* approaches 1 *-m*. Setting RCP = 0 results in two solutions: *U* = 1 -2*m* and *U* = 0. These solutions are congruent with the boundaries that define the space of (*m, U*) as given in “Ratio of Concordance Preference is Defined for All Possible Configurations of Methylation at Symmetric Nucleotide Motifs” of the main text, and in Fig 1.

### S2 Text: Comparing RCP Values and Hemi-Preference Ratios from HMM

“Hemi-preference ratio”, a parameter inferred under our earlier analysis with a hidden-Markov model (HMM) [24], evaluates the preference of a given DNA methyltransferase for acting at hemimethylated as compared to unmethylated dyads [11], and thereby measures its preference for creating concordant dyads. Fu *et al.* [24, 28] calculated this ratio for DNMT1, a mammalian maintenance methyltransferase. Because RCP measures the concordance preference of the entire ensemble of enzymes that give rise to methylation patterns, the hemi-preference ratio of a given enzyme and the RCP value inferred from the same data set are expected to have good agreement if that enzyme is the primary actor. The congruence between these two metrics is expected to decline with increasing contributions from other enzymes.

Three of the four data sets analyzed previously under HMM showed very good agreement between the RCP values we infer here and the hemi-preference ratios previously inferred for the maintenance methyltransferase DNMT1: 58.0 vs. 58 for *FMR1*, 13.3 vs. 15 for *G6PD*, and 89.1 vs. 94 for *LEP* [24] (Table S1). The close correspondence between these values indicates that for these loci in leukocytes, methylation dynamics are driven primarily by conservative, maintenance-type processes such as accomplished by DNMT1, and that neither active demethylation nor *de novo* processes have a substantial role. Furthermore, when there is such a correspondence, RCP strongly suggests that the mechanistic assumptions made for the enzymatic model hold for that data set.

A large discrepancy, on the other hand, may suggest shortcomings of the mechanistic model. The fourth data set that had been analyzed under HMM, *LEP* in human adipose tissue, had an inferred RCP value of 34, well within the range of RCPs inferred for other data sets from single-copy loci including *LEP* in leukocytes (Fig 1a; Table S1). This RCP value was, however, more than eighteen-fold lower than the DNMT1 hemi-preference ratio estimate of 628 that we obtained under the earlier HMM approach (Table S1). What might account for this discrepancy? A hemi-preference ratio of 628 is unrealistically high, even for a maintenance enzyme, compared to the hemi-preference ratios inferred in other data sets, including those from methylation patterns established by DNMT1 *in vitro* [11]. This could reflect the inability of the HMM to yield a reasonable estimate for a data set impacted by demethylation, a process that was not considered in the HMM design. This lack of correspondence between HMM and RCP estimates is consistent with the possibility that active removal of methylation has a heightened role at loci with temporally variable transcription levels, and may reflect the role of the *LEP* locus as a sentinel of adipose but not blood [29].

### S3 Text: Assessing the Potential Impact of Bisulfite-Conversion Errors

Two classes of error that occur during bisulfite conversion can lead to misinterpretation of cytosine methylation states. Failure to convert unmethylated cytosines to uracil occurs at rate *b*, and can result in overestimation of methylation frequency. Inappropriate conversion of methylated cytosines to thymine, first noted by Shiraishi and Hayatsu [50], occurs at rate *c*, and can result in underestimation of methylation frequency. We have reported previously that the rates of failed and inappropriate conversion depend strongly on the chemical and thermal conditions of bisulfite conversion [49]. In particular, we found that bisulfite treatment prolonged beyond that required to attain complete or nearly complete conversion of unmethylated cytosines can yield high rates of inappropriate conversion. Historically, conversion protocols have been designed to minimize the failed-conversion rate, with little or no attention to the rate of inappropriate conversion events. Thus, while both classes of error can alter parameters used to infer RCP — the methylation frequency, *m*, and the unmethylated dyad frequency, *U* — errors arising through inappropriate conversion are likely of more substantial impact.

How severely can conversion error impact RCP? And to what extent does its potential impact depend on the true methylation frequency of the target sequences? Densely methylated sequences contain a large number of fully methylated dyads, such that the most likely conversion error is inappropriate conversion yielding apparent hemimethylated dyads. Such events elevate the apparent level of discordance in a given data set and artifactually depress the inferred RCP values. To quantify this potential impact, we calculated how RCP would be altered by a single artifactual hemimethylated dyad introduced by inappropriate conversion, assuming that in its true state the relevant data set had one unmethylated dyad, one hemimethylated dyad, and 98 methylated dyads. The inappropriate conversion of one cytosine among 98 dyads whose true state is fully methylated is close to the number expected for an inappropriate conversion rate of about 0.5%, a conservative estimate of this error rate [49]. For this hypothetical data set, the introduction of a lone artefactual hemimethylated dyad by an inappropriate conversion event reduces RCP from about 20 to about 10, i.e., by a factor of 2. Thus, in the absence of mathematical corrections of the sort implemented here, even very low levels of inappropriate conversion can severely impact inference of RCP.

Sparsely methylated sequences contain a large number of unmethylated cytosines that are potential targets for failed conversion. We calculated that for such sequences the impact of conversion error depends on whether most unmethylated cytosines are in hemimethylated or in unmethylated dyads. For example, when most unmethylated cytosines in a sparsely methylated sequence occur in hemimethylated dyads, the level of discordance is already high, such that the addition of an artifactual hemimethylated dyad by failed conversion only slightly decreases RCP. By contrast, when most unmethylated cytosines are in unmethylated dyads, production of an artifactual hemimethyated dyad by failed conversion reduces RCP by a factor of two, as illustrated in the previous paragraph. Mathematical correction for conversion error is therefore critical, not only because error has potentially large impacts on RCP values, but also because the magnitude of these impacts differs among data sets, with the potential either to magnify or to dampen variation among them.

### S4 Text: Mathematical Correction for Bisulfite-Conversion Error

To account for conversion errors occurring at known rates, we express the observed dyad frequencies, *M*_*fobs*_, *H*_*fobs*_, and *U*_*fobs*_, as functions of the true dyad frequencies *M*_*t*_, *H*_*t*_, and *U*_*t*_, that would have been observed had conversion error not occurred:

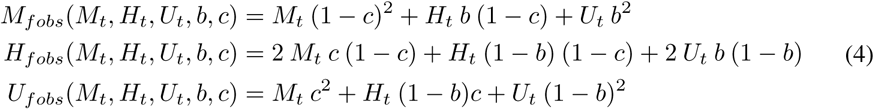

In cases where the mathematical correction yielded negative counts for one or two dyad types, the negative counts were redistributed to the remaining dyad types in proportion to the original dyad counts, such that no dyad counts were negative after correction.

For each of the data sets reported here, DNAs were converted under one of two conditions: low molarity-temperature (LowMT), with failed-conversion rate 0.0030 and inappropriate conversion rate 0.031, and high molarity-temperature (HighMT), with failed-conversion rate 0.0086 and inappropriate-conversion rate 0.017 [49]. Error rates used in analysis of published data from other groups are given in Table S4 (Arand *et al.* [14, 17]) and Table S5 (Zhao *et al.* [15]). Using these rates, observed dyad frequencies, and Equation 4, true dyad frequencies can be inferred.

### S5 Text: Assessment of Heterogeneity Among Replicates and Quasi-Replica

To ask about possible heterogeneity among RCP values inferred for replicates — for example, for individual developmental stages as investigated by Arand *et al.* [17] — and for quasi-replicates — for example, for multiple cell lines at similar cell-culture and differentiation conditions — we utilized pairwise, two-tailed permutation tests.

These calculations revealed an intriguing pattern. There was no evidence of heterogeneity among samples of cultured human ES and iPS cells (*p >* 0.29, pairwise two-tailed PTs for the six cultured stem cell lines at the *L1* locus; *p*-values summarized in Table S6; Fig 3). By contrast, several pairwise comparisons of mouse embryonic cells sampled independently for a given developmental stage yielded evidence of significant differences in RCP (*p*-values summarized in Table S6). Notably, however, most of the significant heterogeneity observed was for cells in pronuclear stages, at which data collection at identical stages is made difficult by rapid developmental transitions. In contrast, there was evidence of only limited heterogeneity among totipotent and pluripotent cells between the 2-cell stage and the 3.5-dpc stage. Similarly, when experimental interventions were taken to prevent DNA replication or methylation (i.e. aphidicolin treatment at PN4/5 stages and SAMase treatment just after fertilization and before the first round of DNA replication), no evidence of heterogeneity was observed. Cell culture techniques are focused on minimizing opportunities for developmental differences to arise among cells, and could account for the observation of no heterogeneity among the cultured human ES and iPS cells.

Given the low levels of heterogeneity observed and the fact that some level of heterogeneity is inevitable in cells undergoing rapid epigenetic transitions, we pooled several sets of replicates for analysis, as described in S6 Text (Fig 6).

### S6 Text: Pooling Data Sets Across Replicates

In some instances, multiple independent replicates were collected and used to assess RCP for cells under a given biological condition — for example, multiple embryos at a given pronuclear stage (Fig 6). In these cases, the sequence-level dyad counts of individual replicate data sets were corrected for conversion errors independently before the replicate data sets were pooled.

When using likelihood methods to quantify and to assess pooled data sets – thus assuming independence of dyads (S8 Text) – we explicitly allow *m* to vary across replicates while estimating a single *RCP* value in the likelihood-maximization process. When using bootstrap methods (S7 Text), as we do whenever individual-molecule data are available, we do not explicitly account for the possibility that *m* may vary across the replicate data sets. This may result in a slight enlargement of confidence intervals in the bootstrap process.

### S7 Text: Inferring and Comparing RCP Without Assuming Independent Sampling of Dyads

CpG dyads typically are not sampled individually, but instead as members of sequence reads that can contain from a few to many neighboring dyads. Often, there is correlation among the methylation states of these neighboring dyads. The processivity of the DNA methyltransferases, especially Dnmt1, is a substantial contributor to these correlations. As the mean number of dyads per read increases, so too does the potential for dyad-dyad correlation to undermine the accuracy of confidence intervals inferred under the assumption of independent sampling of dyads.

Of the three groups of data we analyzed here, one — Zhao *et al.* (2014) [15] — consists of Illumina paired-end reads. These reads span 40 to 60 genomic nucleotides, and so are much shorter than those generated by our methods and those of Arand *et al.*. On average, each of the read pairs in Zhao *et al.* provides methylation data for only 1.2 CpG dyads. As this CpG count is only slightly greater than the condition of 1 CpG dyad per read that would provide for complete independence among sampled dyads, we anticipate that correlation among methylation states of dyads ascertained on a given read by Zhao *et al.* will have only a minor impact on sampling. The subset of these reads that derive from CpG-rich CpG Islands do contain more CpGs, likely up to 3 [51]. However, this sequence class contributes only 1.2% of CpG dyads in the “All” CpG dataset. Reads for our data have as many as 22 CpG dyads; mean dyad counts for data from Arand et al. [14, 17] are intermediate between our data and those of Zhao *et al.* (2014) [15].

For our own data and those of Arand *et al.* [14, 17], we used an approach that, at slightly higher computational cost, models and seeks to account for the potential impacts of dyad-dyad correlations. Our data yielded moderately larger confidence intervals under the bootstrapping approach as compared to under the likelihood approach with the assumption of independent sampling of dyads. By contrast, the data from Arand *et al.* [14, 17] yielded almost identical confidence intervals whether without or with the assumption of independence. In view of the even smaller mean number of dyads per read in the data of Zhao *et al.* (2014) [15], we chose to make the assumption that dyads were sampled independently in their data. Although it is possible that two or more reads originated from nearby regions of a single molecule and thus have dyad-dyad dependence, we assumed that the effect of such occurrences, if at all present, is very small, given the large amount of starting material.

The first of the two methods is described in this section, and the second in the next section, S8 Text.

#### Inferring RCP point estimates and confidence intervals.

For our own data and those of Arand *et al.*, we used a bootstrapping approach to model the uncertainty in RCP values introduced by possible within-sequence correlations in methylation states, and to make point estimate and confidence-interval inferences that account for this uncertainty.

For each data set of *n* sequences, we sampled *n* sequences with replacement, *B* = 2,000,000 times. For each of these bootstrapped sets, RCP was calculated by summing the *M*, *H*, and *U* dyad counts for all of the resampled molecules and using Equation 3. We inferred the true distribution of RCP for a given observed data set from the distribution of these *B* bootstrapped RCP values. It was clear from the resulting distributions that RCP is a biased estimator, as many of these distributions had longer right tails than left.

The simplest, and potentially misleading, approach for inference of point estimates and construction of confidence intervals from bootstrap distributions is to assume normality, and then to exclude right and left tails at the intended level of confidence. Efron and DiCiccio (1996) [52] commented that, for biased estimators, this approach can lead to inference of inappropriately exclusive limits at the long-tailed end of the distribution, and inappropriately inclusive limits at the short-tailed side.

To address this problem, we applied Efron and Diccicio’s “bias-corrected and accelerated” (BCa) method. Under the BCa method, the cumulative-density function observed for the distribution of bootstrap replicates is compared to that expected under normality. Bias-corrected point estimates are then inferred as the 50th-percentile values in the BCa-corrected distributions. Similarly, critical points for intervals of a given confidence level are inferred from the values at the relevant percentiles of the distribution of bootstrapped values.

Because methylation states are predicted to be correlated across dyads within a molecule, but not across molecules, we resample at the level of molecules. Furthermore, as noted above, our molecular-barcoding procedures enable exclusion of redundant reads, such that each methylation pattern in our resulting data set is known to derive from a unique molecule in the original sample. Moreover, it appears that molecules are sampled without bias due to methylation: in all eight data sets from murine DNA methyltransferase knockout lines (Fig 4) and all six data sets from wildtype human cells (Fig 3) there was no evidence of correlation between methylation frequency and RCP. Thus, it is reasonable to consider sampled molecules as independent and identically distributed draws from a population.

For each sampled molecule we derive a vector of values, (*M*_*i*_*, H*_*i*_*, U*_*i*_*, n*_*i*_), where *n*_*i*_ is the number of dyads. These are the vectors we resample in our procedure. We see that RCP can be written, using this vector, as

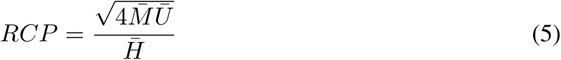

where 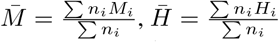 and 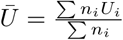. Now considering the vector (*n*_*i*_*M*_*i*_*, n*_*i*_*H*_*i*_*, n*_*i*_*U*_*i*_*, n*_*i*_) and, noting that the ratio of sums can be written as a ratio of means, we see that this falls directly under the “smooth function of means” framework introduced in [53] for the standard bootstrap and applied in [54] to the BCa bootstrap. From results in Section 6 of [54], we conclude that our bootstrap procedure gives approximate confidence intervals with asympotically correct coverage.

#### Assessing whether a data set has RCP value greater than 1.

If methyl groups are placed completely at random — that is, with preference for neither concordance nor discordance — RCP is expected to be 1. In a previous report, Shipony *et al.* [16] interpreted their data to indicate that methyl groups are, indeed, placed essentially at random in undifferentiated cells. As there is very little evidence in any of our data sets to indicate possible preference for discordance, we opted to perform a one-tailed test, asking for the probability that our data sets do not have RCP values greater than 1.

To do so, we first calculated RCP point estimates for 200,000 bootstrap replicates and performed the BCa correction, using the method described above, and then calculated the approximate *p*-value as the fraction of those point estimates that were less than or equal to RCP of 1. For example, from the finding that only 20 of the 200,000 bootstrap replicates yielded an RCP point estimate less than 1, we would conclude that RCP is significantly greater than 1 with an approximate *p*-value of 0.0001. Note that we shifted from the 2,000,000 bootstrap draws noted above to the 200,000 reported here upon finding only trivial differences between *p*-values derived under these two approaches.

#### Assessing whether RCP values differ significantly between data sets.

A key goal of our study is to assess possible evidence for RCP differences between data sets. For example, we ask whether RCP for a given cell type differs between samples collected under differentiating as compared to non-differentiating conditions.

To compare two data sets that can be modeled by the same null distribution, as is the case for most comparisons we make here, we performed a permutation test (PT). For comparisons in which a shared null distribution cannot be established (such as would be the case if the two data sets were from different loci and thus had different numbers and locations of dyads), we used a bootstrap test (BT). We describe these two methods below.

To compute the significance of observed differences between RCP values for Data Sample A and Sample B, which can share a null distribution, we used permutation to compute the null distribution of ordered differences (for example, RCP(Sample A)-RCP(Sample B)) expected under the null assumption that the sequences in the two sets were drawn from a single population. To do this, we pooled sequences from Sample A and Sample B and then repeatedly drew from that pool, without replacement, to generate, at random, versions of Sample A and Sample B with sequence counts equal to those of the observed data. For each pair of randomly generated sets, we calculated the difference between their RCP point estimates (*?*^*\**^), and obtained the distribution of RCP-difference values under the null hypothesis. For one-tailed tests, for example, to ask whether one data set has RCP value greater than the RCP value for another, we took the proportion of the distribution greater than the difference of the point estimates 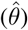 as the *p* value. For two-tailed/equal-tailed tests, for example, to ask whether RCP values for two data sets differ significantly, we first calculated the proportion of permuted differences smaller than 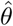. We then calculated the proportion of permuted differences greater than 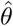. We took the twice the lesser of these two values as the *p*-value for observing a difference this great in the event that the sequences in the two data sets were, in reality, drawn from the same distribution.

To compute the significance of observed differences between RCP values for two samples with different generating distributions, we used a bootstrap-based comparison test. Instead of pooling the data sets to form a single population of molecules, we bootstrapped a single RCP value from each of the two separate populations of molecules and computed the difference. We repeatedly sampled this difference to draw a bootstrap distribution of the difference in RCP values. For one-tailed tests, with which we examine directional differences, we determined the *p*-value as the proportion of bootstrap-difference samples to the left of 0. For two-tailed tests, with which we can detect differences in any direction, we determined the *p*-value as twice the smaller proportion of the bootstrap difference samples on either side of 0.

#### Defining the approximate 95% confidence region for a data set in the (*m, U*) space, without assuming independent sampling of dyads.

Using the methods described above, we generated 2,000,000 bootstrap samples of each data set. Instead of estimating the RCP value for each of the bootstrap samples, we calculated *m* and *U*. We constructed the two-dimensional confidence region for the two parameters for plotting using `ci2d` function in the `gplots` R package.

### S8 Text: Inferring and Comparing RCP With Assuming Independent Sampling of Dyads

#### Calculating the likelihood of proposed true dyad frequencies, *M*_*t*_, *H*_*t*_, and *U*_*t*_, given the observed dyad counts, *M*_*c obs*_, *H*_*c obs*_, and *U*_*c obs*_

Failed- and inappropriate-conversion events create observed dyad frequencies that differ from true dyad frequencies. If the rates of these two types of error are known, the likelihood of a set of proposed true frequencies — *M*_*t*_, *H*_*t*_, and *U*_*t*_ — can be calculated as follows, given the observed dyad counts — *M*_*c obs*_, *H*_*c obs*_, and *U*_*c obs*_ — and Equation (4). *M*_*fobs*_, *H*_*fobs*_, and *U*_*fobs*_, which indicate the observed frequencies of the dyads, can be easily calculated from the dyad counts.

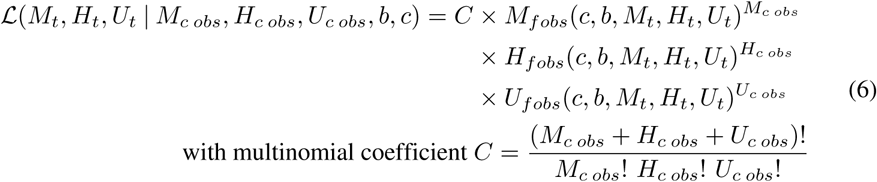

#### Inferring RCP point estimates and confidence intervals

The RCP point estimate of a data set is calculated directly from the conversion-error-corrected observed dyad frequencies. We determine the approximate 95% confidence interval for RCP as the interval that includes all values of RCP for which the natural log likelihood lies within 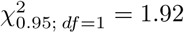 units of the maximum natural log likelihood point estimate [55].

Although bias is just as much a concern with the assumption of independent sampling of dyads, we did not perform a bias correction for the data from Zhao *et al.* [15], because without bootstrapping, we lacked a simple method for bias estimation. Nonetheless, our analyses of other data sets suggest that most of these samples are likely not severely affected by bias. From bootstrapping of smaller data sets collected by our lab and by Arand *et al.* [14, 17], we have observed that the asymmetry in the distribution is small in the lower ranges of RCP (1*∼*10), but greater as RCP increases. Therefore, bias is likely to be small for most samples presented by Zhao *et al.*, for which RCP point estimates rarely exceed 10. For data sets presented by Zhao *et al.* that yielded RCP estimates greater than 10, biases are unlikely to affect the general conclusion of methylation behavior characterized by strong preference for concordance.

#### Assessing whether RCP values differ significantly between two data sets

To compare the RCP values between two data sets while assuming independence among all dyads, we can use a likelihood approach, comparing a model in which two true RCP values are required to described the two data sets to an alternate model in which both data sets can be explained by a single RCP value. We implemented that test as follows:

Solving for *U* in Equation 3 gives:

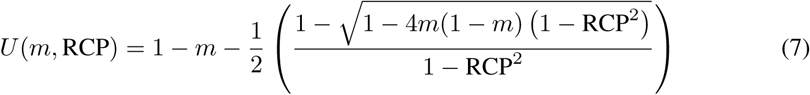

Using this, we can also define *M* (*m,* RCP) and *H*(*m,* RCP). Modifying Equation 4, we can derive the expressions for *M*_*fobs*_(*c, b, m,* RCP), *H*_*fobs*_(*c, b, m,* RCP), and *U*_*fobs*_(*c, b, m,* RCP). We then can rewrite Equation 6, such that the parameters are *c*, *b*, *m*, and RCP:

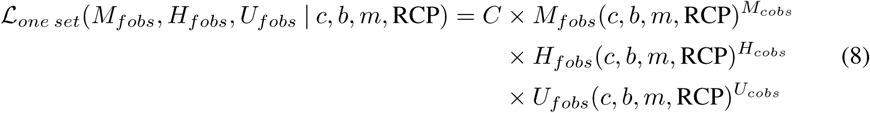

We then employ a likelihood-ratio test to quantify the fit of an alternative model relative to the null. In the null model, which has three variable parameters, 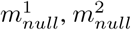, and RCP_*null*_, one value of RCP explains both data sets. Using Equation 8:

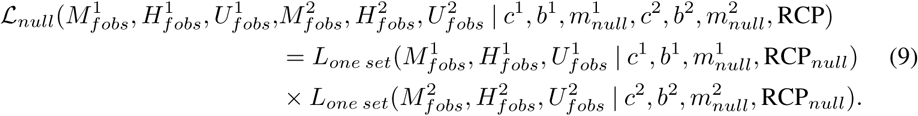

The alternate model, which has four variable parameters, 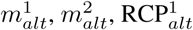, and 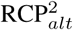, has two values of RCP, one for each data set.

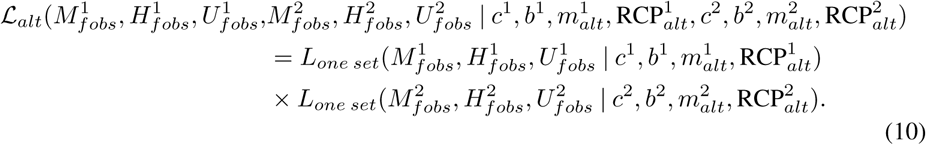

Computing the ratio of the maximum likelihoods for the null and alternate models, we can calculate the test statistic, *D*:

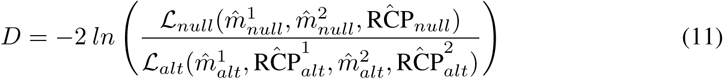

Under the assumption of large sample of dyads, *D* is approximately *χ*^2^ distributed with 1 degree of freedom.

#### Assessing whether a data set has RCP value greater than 1.

We again take a likelihood-based approach as we did in the section above. Here, the null model states that the system operates under the specified RCP value. The alternate model states that the system operates under another RCP value.

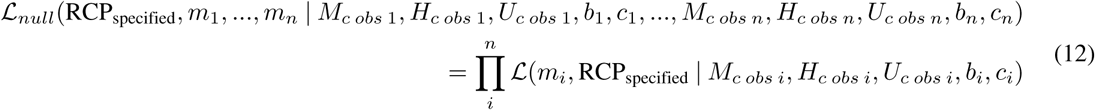

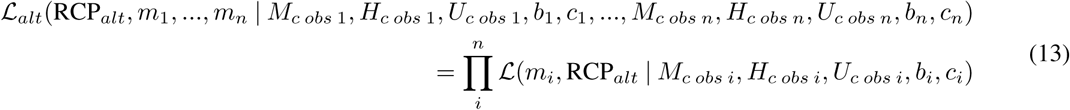

We treat the two likelihood functions differently in maximizing them. For the alternate model, both *n* values of *m* and RCP_*alt*_ are parameters for maximization. For the null model, the RCP value is specified and thus fixed; only the *n* values of *m* are parameters for maximization.

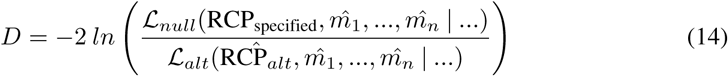

Under the assumption of a large sample of dyads, *D* is approximately *χ*^2^ distributed with 1 degree of freedom.

#### Defining the approximate 95% confidence region for a data set in the (*m, U*) space

We determine the approximate 95% confidence region in the space of two parameters — here *m* and *U* — as the region that includes all proposed pairs of parameter values for which the natural log likelihood lies within 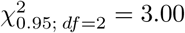 units of the maximum natural log likelihood point estimate [55].

### S9 Text: Could Spontaneous Differentiation of a Subset of ES and iPS Cells Substantially Influence the Inference of RCP?

One possible explanation for the inference of conservative processes operating in cultured, undifferentiated cells is that these cells may in reality be a mixture of differentiated and undifferentiated cells. We calculated how large the differentiated subpopulation would need to be under this scenario to yield the observed RCP values for the ES and iPS cells, given that the subpopulation operated with the RCP inferred for the corresponding differentiated cells. The remainder of the population was assumed to operate at RCP of 1, per the alternate hypothesis. We allowed specification of *m* for each of the two populations.

Let *p*_1_ and *p*_2_ represent the proportions of the two putative subpopulations that we wish to estimate, such that *p*_1_ + *p*_2_ = 1. We start with RCP and *m* for each of the two subpopulations, which we denote by RCP_1_, RCP_2_, *m*_1_, and *m*_2_. We can find *U*_1_ and *U*_2_ using the RCP and *m* values. We have the following set of equations:

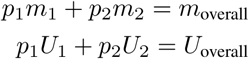

Using these equations and the observed overall RCP value, and the expression for RCP given *m* and *U* (Equation (3)), we can find *p*_1_ and *p*_2_.

Assuming that *m* of the undifferentiated subpopulation is at least 0.2, we found that over half of the cells would need to be differentiated if methylation in the remaining undifferentiated cells operates at RCP of 1 (calculated using data sets presented in Fig 5a). Morphological inspection of the cultured human stem cells did not suggest the presence of such a substantial subpopulation of differentiated cells. Moreover, comparison of RCP for totipotent and pluripotent stem cells from murine embryos revealed values very similar to those inferred for all cultured ES and iPS cell lines that we examined, corroborating the interpretation that preference for concordance is present in stem cells.

### S10 Text: Comparing RCPs of *Dnmt3*-Knockout Lines with Those of Wildtype Lines

The relative contributions of conservative processes are expected to be more substantial when the fraction of methylation events achieved through maintenance-type activity is elevated through loss of one or both of the *de novo* enzymes. Indeed, at the *Lep* locus, we inferred RCP values of 8.11 for *Dmnt3a* KO cells, a value significantly higher than that for wildtype cells (*p* = 0.046, two-tailed PT; Fig S2a; Table S3). Although the point estimate was higher for *Dmnt3b* KO cells as well, the difference was not significant (*p* = 0.11, two-tailed PT).

Examination of data from Arand *et al.* [14] for a broader set of loci in cell lines with knockouts at one or both of the Dnmt3s, however, revealed a more complex role for these enzymes in shaping methylation concordance. Overall methylation levels were diminished in the absence of the *de novo* methyltransferases 3a and 3b, as expected (Fig S2b-i, Table S3); RCP values increased as expected for several, though not all, loci. Results for cells that lacked only one of the two DNA methyltransferases were even more variable across loci (Fig S2, Table S3). In some cases, loss of a single Dnmt3 enzyme had the predicted impact of increasing RCP (*L1* in 3a KO, *p <* 10^-16^, Fig S2h; *L1* in 3b KO, *p* = 0.002, Fig S2h; *IAP* in 3a KO, *p* = 0.002, Fig S2g; *IAP* in 3b KO, *p* = 0.029, Fig S2g; two-tailed PTs), as was observed for *Lep* in our data. For one locus, *B1*, knockout of both of the Dnmt3s resulted in significantly increased RCP (*p <* 10^-16^, two-tailed PT), though neither of the single knockouts did (Fig S2f). Most surprisingly, for two loci, loss of either of the Dnmt3s decreased RCP (*Igf2*, *p <* 0.016, Fig S2c; *Igf2*, *p <* 10^-16^, Fig S2c; two-tailed PTs). These counterintuitive results likely reflect variation across loci in the roles of the individual DNA methlytransferases — and possibly the demethylation machinery — in shaping overall methylation levels for various loci and categories of genic elements [56].

